# Tilted striatofugal balance and beneficial effects of facilitating mGlu4 receptor activity in the Fmr1^-/-^ mouse model of Fragile X Syndrome

**DOI:** 10.64898/2026.01.12.699042

**Authors:** Mathieu Fonteneau, Claire Terrier, Florian Bolot, Fani Pantouli, Fanny Malhaire, Roser Borràs-Tuduri, Isabelle McCort-Tranchepain, Alexis Bailey, Francine Acher, Amadeu Llebaria, Cyril Goudet, Jérôme A.J. Becker, Julie Le Merrer

## Abstract

Fragile X Syndrome (FXS) is the leading monogenic cause of autism spectrum disorder (ASD). To date, no approved pharmacological treatment alleviates social impairments in patients with FXS. Since D1 and D2 dopamine receptor-expressing striatal projection neurons (SPNs) were shown to regulate ASD-sensitive behaviors, we explored whether the balance of activity between D1- and D2-SPNs would be biased in the *Fmr1^-/-^* mouse model of FXS. We evaluated striatal function in *Fmr1*^-/-^ and *Fmr1*^+/+^ mice under pharmacological challenge and performed RNAscope® *in situ* hybridization following social interaction to assess the activity of SPNs in the nucleus accumbens (NAc) and dorsal striatum (DS). We evidenced a decrease in D1-SPN activity, biasing the D1/D2-SPN balance towards an excessive weight of D2-SPN outputs. We then evaluated the effects of compounds that repress D2-SPN activity on behavioral impairments in *Fmr1*^-/-^ mice. Systemically facilitating mGlu4 or blocking A2A receptor activity relieved behavioral deficits in this model. Finally, we tested the hypothesis that a tilt of the D1/D2-SPN balance in *Fmr1*^-/-^ mice may contribute to their social deficit by facilitating mGlu4 activity directly in the projection site of NAc D2-SPNs. Social interaction in *Fmr1*^-/-^ mice was fully rescued by photopharmacological activation of mGlu4 in the ventral pallidum (VP), where NAc D2-SPNs project. This result supports our hypothesis of excessive D2-SPN outputs and highlights the contribution of the VP in controlling social behavior. In conclusion, pharmacological compounds that repress D2-SPN activity demonstrate a promising therapeutic potential to relieve ASD-like deficits in FXS.

## 1. Introduction

Autism spectrum disorders (ASD) are complex neurodevelopmental disorders whose diagnosis is reached in presence of impaired social communication and interaction together with a restricted, repetitive repertoire of behaviors, interests and activities [1]. The etiopathological mechanisms underlying ASD remain essentially unknown, although a clear genetic contribution has been identified [2,3]. Fragile X syndrome (FXS) is the most prevalent monogenic cause of ASD, accounting for 1 to 6 % of total cases [4–6]. FXS results from CGG repeat expansion (>200) in the *Fmr1* gene, impeding the transcription of the Fragile X Messenger Ribonucleoprotein (FMRP). About 50% of males and 20% of females with FXS meet the diagnosis for ASD; other behavioral and neurologic challenges include intellectual disability, generalized anxiety, hyperactivity and epilepsy [4,5,7]. Current treatments target comorbid symptoms such as anxiety, sleep problems or epilepsy [5,8] but fail to relieve cognitive impairment or core social deficits of ASD. Preclinical evidence of cellular and behavioral benefits of mGlu5 inhibitors [9,10] raised major hopes for FXS treatment, but these compounds disappointingly failed in clinical trials [11,12].

Complex neuropsychiatric profile in FXS points towards widespread alterations of the central nervous system. Indeed, as an RNA-binding protein, FMRP regulates the expression of multiple key components of synaptic function throughout the brain [13–15]. The high prevalence of learning disabilities and cognitive deficits in this syndrome have led to an extensive investigation of cortical [16–18] and hippocampal [19,20] dysfunction in this disorder. However, clinical and preclinical evidence for motor impairment, behavioral inflexibility, reward and social deficits in FXS [4,21,22] point towards altered striatal function, as proposed for ASD [23,24]. Within the striatum, the nucleus accumbens (NAc), a hub structure of the reward circuitry [25–27] that plays a unique role in reward processing and goal-directed behavior [25,28], has received little attention in FXS. This region plays a key role in modulating social and repetitive behaviors [29–31], questioning a potential contribution of NAc dysfunction to ASD-like symptoms in FXS.

GABAergic striatal projection neurons (SPNs) represent the main neuronal population in the NAc, and segregates into dopamine D1 receptor (D1) and D2 receptor (D2)-expressing subtypes. A finely tuned balance of activity between D1- and D2-SPNs in the NAc drives approach and avoidance behaviors [32], including towards congeners [30,33–36]. Convergent evidence points towards altered NAc physiology in the mouse model of FXS, the *Fmr1^-/-^* mice. NAc SPNs indeed display altered synaptic plasticity and dendritic morphology in this model [37,38]. More specifically, the lack of FMRP reshapes the electrophysiological profile of D1- and D2-SPNs in the NAc Core, favoring D2-SPN outputs [39]. Moreover, the functional connectivity of the NAc is modified in a pathway-dependent manner in *Fmr1^-/-^* mice: the synaptic strength of prefrontal cortex (PFC) and basolateral amygdala (BLA) projections to NAc Core D2-SPNs is increased, together with dendritic spine density of NAc Core D2-SPNs [40]. Given the role of NAc D2-SPNs in controlling social behavior [30,35,41], one would consider plausible that excessive activity of NAc D2-SPNs would participate in the ASD-like symptomatology in FXS; as such, pharmacological compounds repressing the activity of this neuronal population should alleviate these symptoms. In support of this hypothesis, facilitating the activity of the metabotropic glutamate receptor 4 (mGlu4), highly expressed on SPN terminals and known to inhibit GABA release D2-SPNs terminals [42–44], or blocking the A2A adenosine receptor, whose expression is highly enriched in D2-SPNs [45,46], were recently found to improve cellular and behavioral features in *Fmr1^-/-^*mice [47,48].

In the present study, we explored striatal-dependent behaviors in *Fmr1^-/-^* mice in response to dopaminergic and opioid compounds, in search of functional evidence of a bias in the D1/D2-SPN balance of activity. Using the RNAscope® in situ hybridization (ISH) technique, we assessed the activity of D1- and D2-SPNs in the ventral (NAc Core and Shell) and dorsal (DS) striatum following a social encounter in *Fmr1*^-/-^ and *Fmr1*^+/+^ mice. Then, we evaluated the therapeutic potential of four compounds known to suppress D2-SPN activity: two positive allosteric modulators (PAMs: VU0155041 and ADX88178) and an orthosteric agonist (LSP4-2022) of mGlu4, as well as an A2A antagonist, istradefylline [49–52]. We examined their ability to alleviate ASD-like symptoms in *Fmr1^-/-^* mice after systemic administration. Finally, we tested whether unilateral photopharmacological stimulation of mGlu4 with optogluram, a photoswitchable mGlu4 PAM, in the projection site of NAc D2-SPNs, the ventral pallidum (VP), would be sufficient to rescue social behavior in *Fmr1^-/-^* mice. Our results confirm the therapeutic promises of pharmacologically repressing D2-SPN activity to relieve ASD-like symptoms in FXS.

## 2. Materials and methods

### 2.1. Animals, housing conditions and breeding procedures

Equivalent numbers of male and female *Fmr1^+/+^* and *Fmr1^-/-^* mice (KO2, generously provided by R. Willemsen, Erasmus University Medical Center, Rotterdam, The Netherlands) were bred in-house on a C57BL/6J background. *Fmr1^+/+^* and *Fmr1^-/-^* pups were bred from homozygous parents, which were bred from heterozygous animals, to prevent genetic drift. Mice in the same cage were of the same genotype: this breeding scheme likely exacerbated behavioral deficits in *Fmr1^-/-^* mice by maintaining them together during early post-natal development [53]. We defined sample size (G*Power 3.1) to ensure sufficient statistical power using ANOVA or Kruskal-Wallis analysis of variance to detect significant effects on our parameters (effect size f=1.80, α=0.05, 0=5, n=8, power=0.96). Mice were weaned at 3-week age. Cages containing *Fmr1^+/+^* or *Fmr1^-/-^* mice (same age and sex) were organized from as many different litters as possible (to limit litter effects) by the staff of the animal facility (blind to experiments) and assigned randomly to a treatment condition (same treatment in the whole cage). We ensured that sex ratio was equivalent between groups, and that mice from different litters met during the direct social interaction test. Except otherwise stated, animals were group-housed and maintained on a 12hr light/dark cycle (lights on at 7:00 AM) at controlled temperature (21±1°C); food and water were available *ad libitum*. Experiments were analyzed blind to genotypes and experimental conditions. All experimental procedures were conducted in accordance with the European Communities Council Directive 2010/63/EU and approved by the Comité d’Éthique en Expérimentation animale Val de Loire (C2EA-19).

### 2.2. Drugs for in vivo experiments

The D1/D5 dopamine receptor agonist SKF81297 (hydrobromide), the D2/D3 dopamine receptor agonist quinpirole (hydrochloride), the non-selective dopamine receptor antagonist haloperidol (hydrochloride), the D1/D5 dopamine receptor antagonist SCH23390 (hydrochloride), the D2/D3 dopamine receptor antagonist sulpiride, the mGlu4 PAM VU0155041 (hydrochloride) and the A2A adenosine receptor antagonist istradefylline were purchased from Tocris (Tocris Bioscience, Bristol, UK). Morphine (hydrochloride) was obtained from Francopia (Antony, France). The mGlu4 PAM ADX88178 was purchased from abcr (Karlsruhe, Germany; ref # AB494741). The mGlu4 orthosteric agonist LSP4-2022 was synthesized in the laboratory of F.A. following previously described procedure [54]. The mGlu4 photoswitchable PAM optogluram was synthesized in the laboratory of A.L. following a previously described procedure [55]. Compounds were dissolved in sterile isotonic saline solution (NaCl 0.9%) except for istradefylline and ADX88178, suspended in 1% carboxymethyl cellulose (CMC, in NaCl 0.9%). Volume of injection (s.c. or i.p.) was 10 ml/kg. Doses refer to salt weight.

### 2.3. RNAscope in situ hybridization

#### 2.3.1. Sacrifice

*Fmr1^+/+^* and *Fmr1^-/-^* mice were tested for direct interaction for 10 min and were sacrificed by cervical dislocation 45 min after the beginning of the test to capture *Fos* mRNA expression induced by the social task.

#### 2.3.2. Brain preparation and RNAscope in-situ hybridization assay

Immediately after the sacrifice, fresh brain tissue was gently frozen in cryomolds with OCT on dry ice and stored at -80°C until sectioning and running RNAscope assay (RNAscope® Multiplex Fluorescent Reagent kit v2, 323100).

Brains were sectioned using a cryostat (Leica CM1860). 16-µm NAc slices were collected on Superfrost Plus slides (Fisher Scientific) and stored at -80°C until performing the RNAscope ISH for *Fos*, *Pdyn* and *Penk* mRNAs. We ran the RNAscope assay according to the user manual for fresh-frozen tissue. On the first day, slices were post-fixed in 4% PFA for 15 min at 4°C and then were dehydrated in successive EtOH baths of increasing concentration (50, 70 and 100% EtOH). Drops of RNAscope hydrogen peroxide were put on dehydrated slides to cover the entire sections for 10 min at room temperature and a hydrophobic barriers surrounded slices were created afterwords. Slides were incubated with the RNAscope Protease IV for 30 min at 40°C and then with a mix of probes targeting *Pdyn* (RNAscope® Probe – Mm-Pdyn-C2, 318771-C2), *Penk* (RNAscope® Probe – Mm-Penk-C3, 318761-C3) and *Fos* (RNAscope® Probe – Mm-Fos-C1, 316921-C1) mRNAs for 2h at 40°C. Slides were stored in 5X SSC overnight at room temperature. On the second day, the signal probes were amplified by using the AMP1, 2 and 3 solutions respectively and were developed by using the HRP-C1, C2, and C3 and the fluorophore solutions for labelling the C1 probe in green (Opal 570 reagent pack, FP1488001KT), the C2 probe in orange (Opal 520 reagent pack, FP1487001KT) and the C3 probe in far red (Opal 690 reagent pack, FP1497001KT). Slides were washed with 1X Wash buffer between each steps described above. In the end, nuclei were labelled by using DAPI solution and slides were mounted with ProLong Gold antifade mountant and stored in the dark at 2-8°C.

#### 2.3.3. Image acquisition

Sections of NAc Core, NAc Shell, lateral and medial DS were imaged on Zeiss Imager Z.2 microscope. For each region, four to twenty-four images (900 µm x 700 µm, 40X magnification) from one to three serial sections (left and right hemispheres, NAc AP: +1.54 mm to +1.34 mm; DS AP: +1.34 mm to +0.98 mm) were selected for each mouse. Single-labelled *Pdyn+* or *Penk+* cells and double-labelled *Fos^+^*-*Pdyn^+^* and *Fos^+^*-*Penk^+^* cells were manually counted by two different experimenters (ImageJ v2.14.0). The percentage of double-labelled cells compared to single-labelled ones was calculated for each type of SPN. Then, a ratio was derived from the two percentages.

### 2.4. Real-time quantitative PCR analysis

Brains were removed and placed into a brain matrix (ASI Instruments, Warren, MI, USA). Prefrontal cortex (PFC), caudate putamen (DS), nucleus accumbens (NAc), central amygdala (CeA) and ventral tegmental area/substantia nigra pars compacta (VTA/SNc) were punched out from 1mm-thick slices (see Figure S4A). We evaluated the expression of transcripts coding for metabotropic glutamate receptors present in cerebral tissue: mGlu1-5 and 7-8 (*Grm1-5* and *7-8*), marker genes of D1- and D2-SPNs (*Drd1, Drd2, Penk, Pdyn, Adora2a, Arpp21, Hpca, Nr4a1, Oprm1*) as well as genes coding for the neuropeptide oxytocin and its receptor (*Oxt, Oxtr*). Tissues were immediately frozen on dry ice and kept at -80°C until use. For each structure of interest, genotype and condition, samples were prepared and processed individually (n=11 mice per genotype). RNA was extracted and purified using the Direct-zol RNA MiniPrep kit (Zymo Research, Irvine, USA). cDNA was synthesized using the ProtoScript II Reverse Transcriptase kit (New England BioLabs, Évry-Courcouronnes, France). qRT-PCR was performed in quadruplets on a CFX384 Touch Real-Time PCR Detection System (Biorad, Marnes-la-Coquette, France) using iQ-SYBR Green Supermix (Bio-Rad) kit with 0.25 µl cDNA in a 12 µl final volume in Hard-Shell Thin-Wall 384-Well Skirted PCR Plates (Biorad). Gene-specific primers were designed using Primer3 software to obtain a 100- to 150-bp product and purchased from Sigma-Aldrich (Saint Quentin, France); sequences are displayed in Table S1. Relative expression ratios were normalized to the expression level of β-actin and the 2^−ΔΔCt^ method was applied to evaluate differential expression level.

### 2.5. In cellulo pharmacology

#### 2.5.1. Cell culture

*Fmr1^-/-^* and *Fmr1^+/+^* HEK293 cells (CL0044661816A) were obtained from Edigene (Beijing, China). Cells were cultured in Dulbecco’s modified Eagle medium (DMEM) complemented with 10% fetal bovine serum (Thermo Fisher Scientific). Cells were maintained at 37°C in a humidified atmosphere with 5 % CO2. Cells were transiently transfected with human A2A, mGlu4 or mGlu5 using lipofectamine 2000 (Thermo Fisher Scientific) following the manufacturer’s recommendations. All receptors contained a N-terminal SNAP-tag to determine their cell-surface expression. Following transfection, cells were seeded overnight at 100,000 per well into black, clear-bottom 96-well culture plates (Greiner Bio-One). Cell-surface expression of the receptors and pharmacological experiments were performed 24 h after transfection.

#### 2.5.2. Cell-based pharmacological assays

The activation of A2A and mGlu4 receptors was evaluated by measuring cAMP accumulation, reflecting their coupling to the Gs and Gi/o pathways, respectively. cAMP levels were quantified using assay kits from Cisbio Bioassays (Codolet, France), following the manufacturer’s instructions. mGlu5 receptor activation, which signals through the Gq pathway, was assessed by monitoring intracellular Ca²⁺ release using the fluorescent probe Fluo4-AM (Thermo Fisher Scientific), as previously described [56] . Cells were exposed to increasing concentrations of reference agonists (Tocris Biosciences, Bristol, UK): NECA for A2A receptors, L-AP4 for mGlu4 receptors (in the presence of 300 nM forskolin), and quisqualate for mGlu5 receptors.

#### 2.5.3. Cell-surface expression

Cell surface receptor expression was assessed using SNAP-tagged receptors labeled with Lumi4-Tb. Twenty-four hours after transfection, cells were incubated with SNAP-Lumi4-Tb, washed with Tag-Lite buffer, and the fluorescence signal was subsequently measured (Cisbio Bioassays).

### 2.6. Behavioral experiments

Equivalent numbers of naive male and female animals were used in each group. All animals were aged 8-12 weeks at the beginning of experiments. Acute pharmacological treatment (Figures 1, 6 and S7) and RNAscope experiments (Figure 2) were performed in independent cohorts of naive mice. For each chronic treatment experiment, behavioral tests were performed successively in a dedicated cohort of naive mice (Figures 3-5), except for the istradefylline experiment, for which two cohorts were used (see timeline of experiments in Figure S8). Behavioral testing order was chosen to minimize the incidence of anxiety generated by each test on later assays. Experiments were conducted and analyzed blind to genotype and experimental condition.

**Figure 1.**
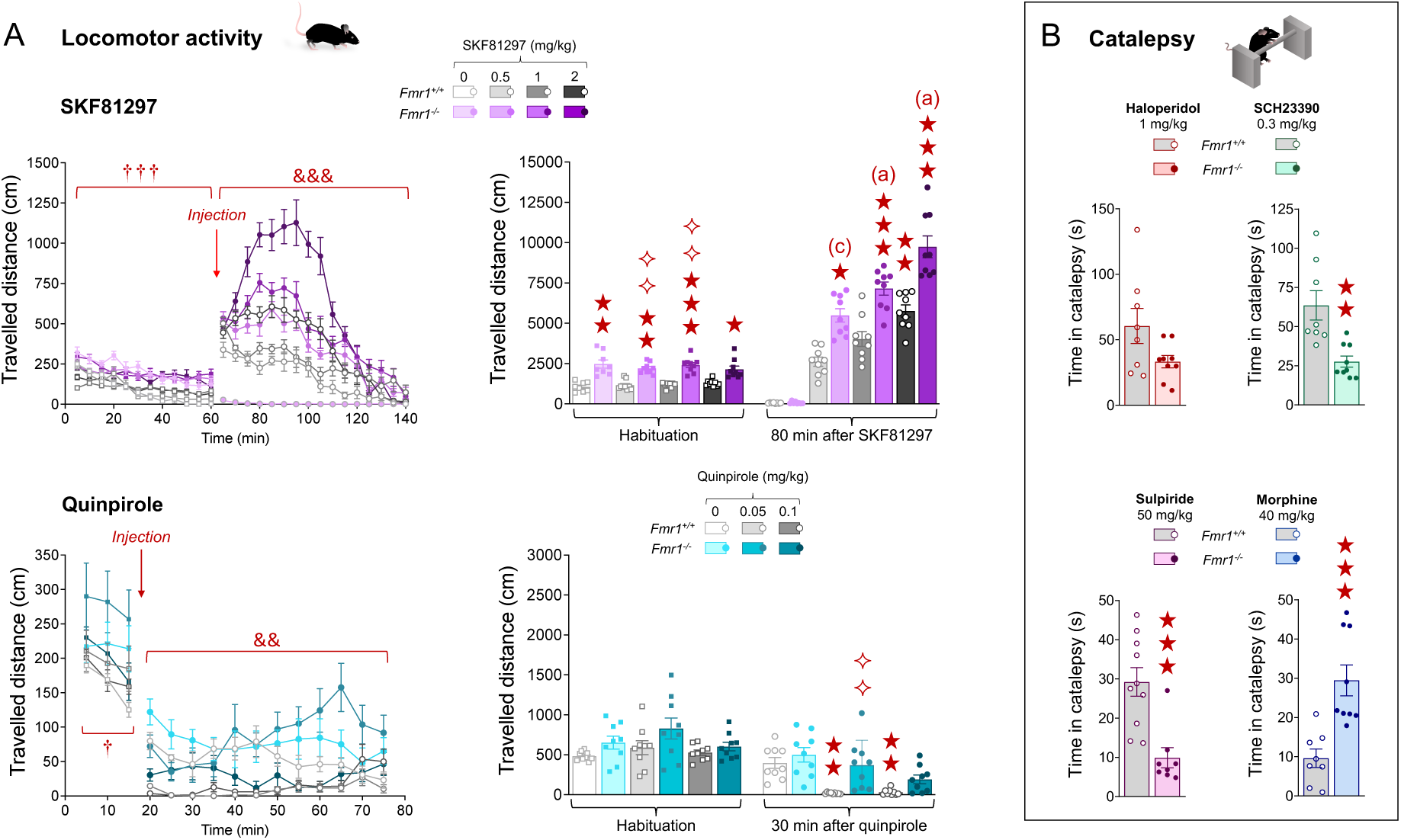
Effects of dopaminergic drugs on locomotor activity and catalepsy in *Fmr1^-/-^*mice. **(A)** Adult *Fmr1^+/+^* and *Fmr1^-/-^* mice received the dopamine D1/D5 receptor agonist SKF81297 (0, 0.5, 1 and 2 mg/kg, s.c.; n=8-9 mice per group) or the D2/D3 receptor agonist quinpirole (0, 0.05 and 0.1 mg/kg, s.c.; n=9 mice per group) after a 60 or 15-min habituation period, respectively. *Fmr1^-/-^* mice were more active than *Fmr1^+/+^* mice during habituation (SKF81297, time course: *G: F_1,46_=88.1, p<0.001*; travelled distance in 80 min: *H_7,70_=52.5, p<0.001*; quinpirole, time course: *G: F_1,48_=6.5, p<0.05;* travelled distance in 30 min: NS). SKF81297 produced higher locomotor stimulation in *Fmr1^-/-^*compared to *Fmr1^+/+^* mice (time course: *G x T x D: F_30,690_=3.1, p<0.001;* travelled distance in 80 min: *H7,70=61.7, p<0.001*). Conversely, quinpirole exhibited a reduced efficacy in decreasing locomotion in *Fmr1^-/-^* mice (time course: *G x T x D: F_22,528_=2.0, p<0.01*; travelled distance in 30 min: *H_5,54_=35.1, p<0.001*). **(B)** Independent groups of adult *Fmr1^+/+^* and *Fmr1^-/-^* mice received the dopaminergic antagonist haloperidol (D1/D2 dopamine receptors, 1 mg/kg), the D1/D5 receptor antagonist SCH23390 (0.3 mg/kg), the D2/D3 receptor antagonist sulpiride (50 mg/kg) 30 min prior testing or the opioid receptor agonist morphine (40 mg/kg) 60 min prior testing. Haloperidol induced a similar cataleptic response in *Fmr1^+/+^* and *Fmr1^-/-^* mice; SCH23390 (*U=4.0, p=0.001*) and sulpiride (*U=4.0, p<0.001*) produced shorter cataleptic responses in *Fmr1^-/-^* mice compared to *Fmr1^+/+^* mice. Conversely, *Fmr1^-/-^*mice treated with morphine spent longer time in catalepsy that *Fmr1^+/+^*mice (*U=2.0, p<0.001*). Results are shown as scatter plots and mean ± SEM. Daggers: genotype effect; ampersands: genotype x time x dose interaction; solid stars: comparison with the vehicle-treated *Fmr1^+/+^*group in locomotor activity experiments (either during habituation or after compound administration), and with the *Fmr1^+/+^* group in catalepsy assays; diamonds: comparison with the *Fmr1^+/+^* group treated by the same dose of treatment. One symbol: p<0.05; two symbols: p<0.01; three symbols: p<0.001. Letters: comparison with vehicle-treated *Fmr1^-/-^* group; (c): p<0.05, (b): p<0.01, (a): p<0.001. Abbreviations: D: dose; G: genotype; NS: not significant; T: time.

**Figure 2.**
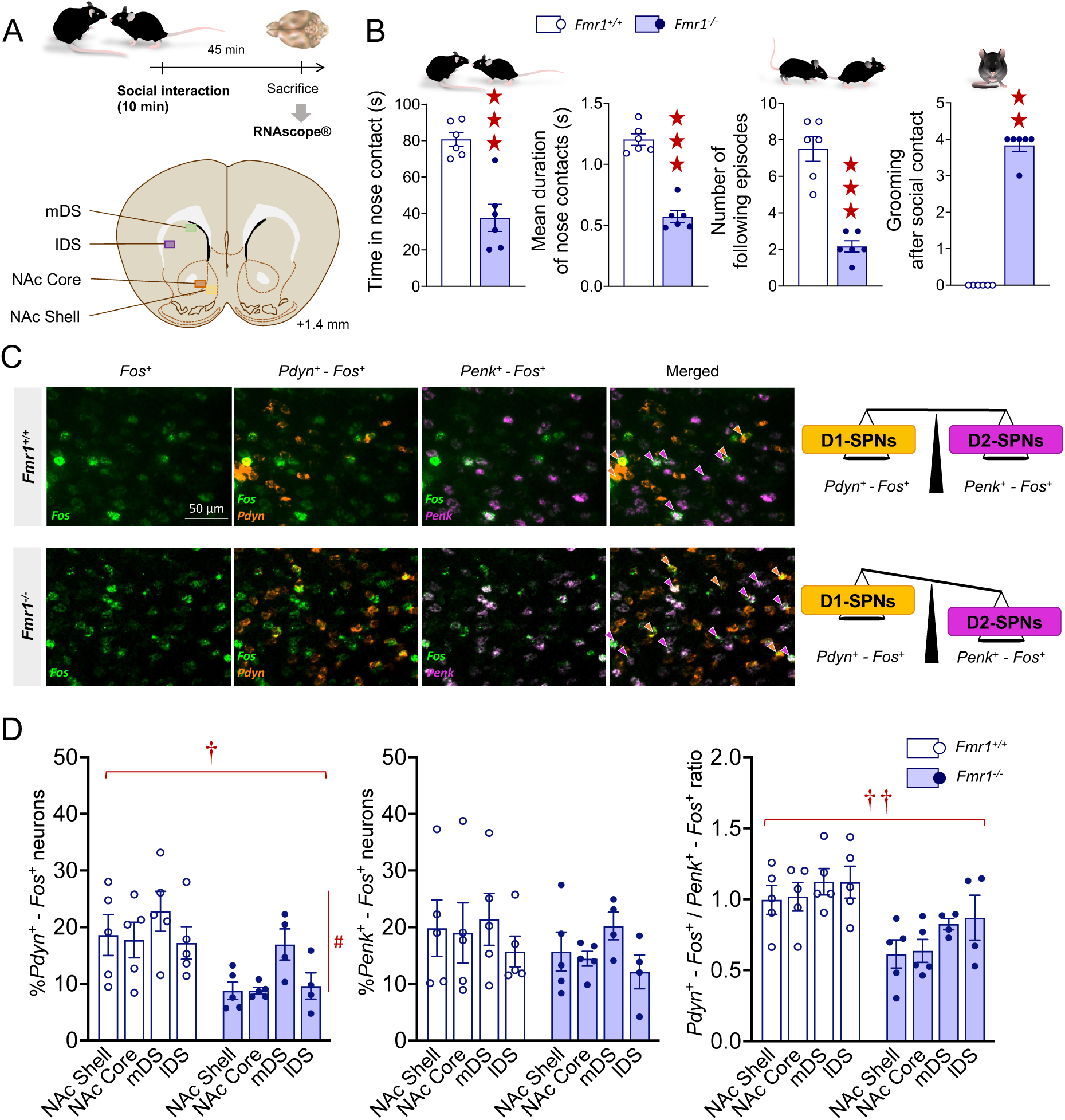
Assessment of *Fos*-positive of D1- and D2-SPNs following direct social interaction in *Fmr1^-/-^*mice using RNAscope® hybridization. (A) Experimental design. Adult *Fmr1^+/+^*and *Fmr1^-/-^* mice (n=6 per group) performed a direct social interaction test (10 min). Mice were euthanized 45 min after the beginning of the interaction test, their brain removed and frozen at -80°C for RNAscope® experiments. **(B)** In the direct social interaction test, *Fmr1^-/-^* mice displayed a severe deficit in social behavior, as evidenced by decreased time spent in (*G: F_1,10_=26.0, p<0.001*) and mean duration of (*G: F_1,10_=89.0, p<0.001*) nose contacts, and number of following episodes (*G: F_1,10_=52.2, p<0.001*), associated with an increase in the number of grooming episodes after social contact (*U=0.0, p<0.01*). More behavioral parameters are available in Figure S3A. **(C)** Representative microphotographs of the NAc Core showing *Fos*-labelled nuclei (green probe), *Pdyn*-labelled neurons (D1-SPNs, orange probe) and *Penk*-labelled neurons (D2-SPNs, red probe) in *Fmr1^+/+^* (upper panel) and *Fmr1^-/-^*(lower panel) mice using the RNAscope® hybridization technique. Magnification: 40x, Scale bar: 50 µm. **(D)** We detected a decreased proportion of *Pdyn*^+^ (D1-SPNs)-*Fos*^+^ neurons in the whole striatum of *Fmr1^-/-^* mice (*G: F_1,10_=9.3, p<0.05*), while the percentage of *Penk^+^* (D2-SPNs)-*Fos^+^* neurons was unaffected. As a consequence of this, the ratio of *Pdyn*^+^-*Fos*^+^ / *Penk^+^*-*Fos^+^* neurons was globally diminished in the striatum of *Fmr1^-/-^*mice (*G: F_1,10_=10.3, p<0.01*). No difference in the total numbers of *Pdyn*^+^ and *Penk^+^* neurons was detected between *Fmr1^+/+^* and *Fmr1^-/-^*mice (Figure S3B). Behavioral and hybridization data are shown as scatter plots and mean ± SEM. Solid stars: comparison with the *Fmr1^+/+^*group; daggers: genotype effect; hashtag: region effect. One symbol: p<0.05; two symbols: p<0.01; three symbols: p<0.001. Abbreviations: G: genotype effect; lDS: dorsal striatum, lateral part, mDS: DS, medial part; NAc Core: core of the nucleus accumbens; NAc Shell: shell of the nucleus accumbens.

**Figure 3.**
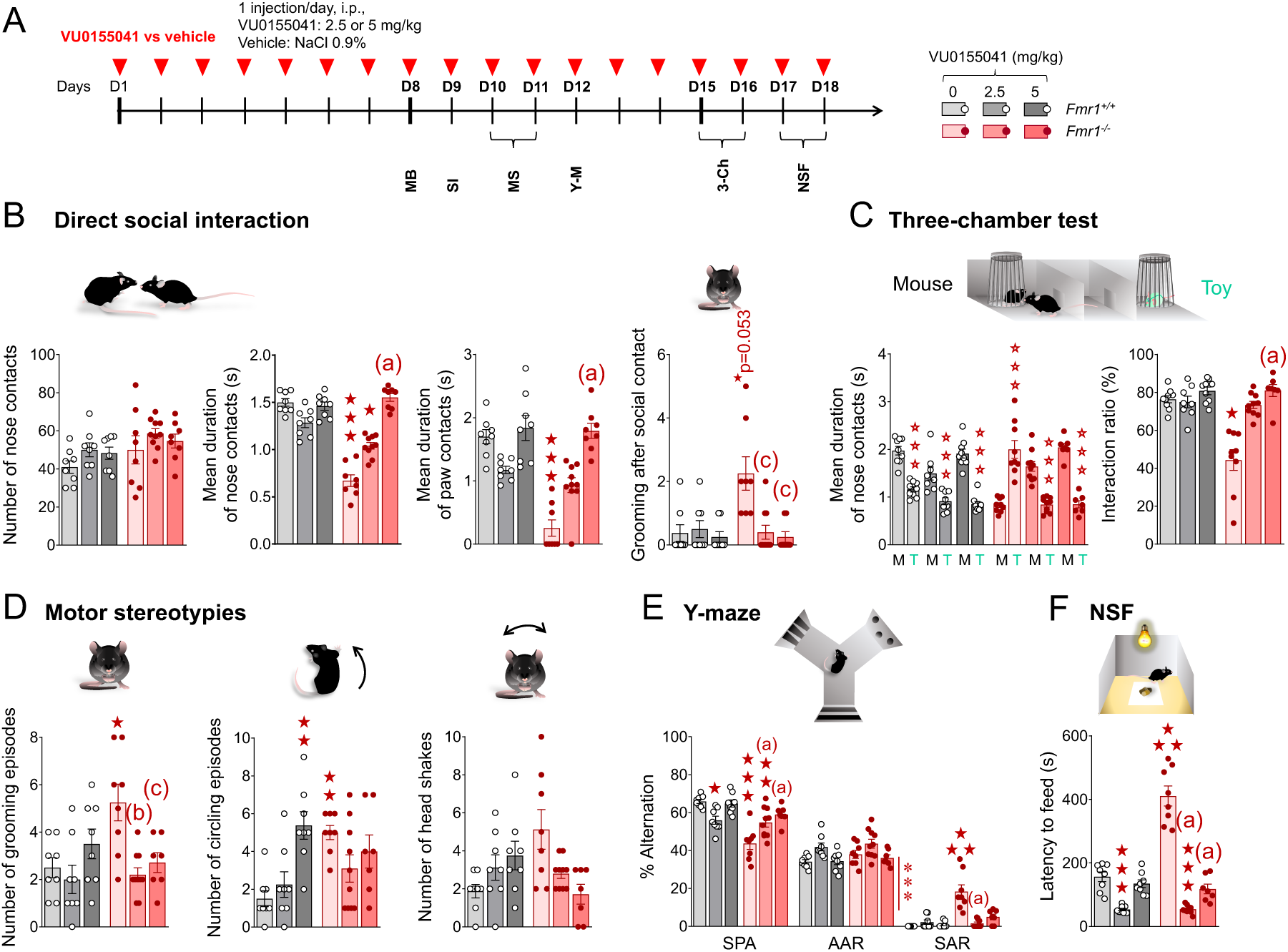
Effects of chronic facilitation of mGlu4 activity on social behavior, motor stereotypies, perseverative behavior and anxiety-like levels in *Fmr1^-/-^*mice. (A) A first cohort of *Fmr1^+/+^* and *Fmr1^-/-^* mice (n=7-10 per group) was injected once daily with either VU0155041 (2.5 or 5 mg/kg, i.p.) or vehicle (NaCl 0.9%) for 18 days. Behavioral testing started on D8. **(B)** In the direct social interaction test, chronic VU0155041 administration dose-dependently normalized nose (*H_5,50_=41.2, p<0.001*) and paw (*H_5,50_=36.4, p<0.001*) contact duration as well as grooming after social contact (*H_5,50_=18.6, p<0.01*) in *Fmr1^-/-^*mice. **(C)** In the three-chamber test, VU0155041 restored the preference for spending longer time (*G x Tr x S: F_2,46_=41.7, p<0.001*) with the mouse versus the toy and preference ratio (*H_5,52_=28.1, p<0.001*) since the 2.5 mg/kg dose. **(D)** Chronic VU0155041 treatment normalized the number of stereotypic grooming (*G x Tr: F_2,43_=3.5, p<0.05*) and circling episodes (*G x Tr: F_2,43_=6.5, p<0.01*) in *Fmr1^-/-^* mice. **(E)** In the Y-maze, VU0155041 restored spontaneous alternation (SPA; *G x Tr: F_2,43_=12.5, p<0.001*) and suppressed perseverative same arm returns (SAR; *H_5,49_=29.5, p<0.001*). **(F)** In the novelty-suppressed feeding test, both doses of VU0155041 fully normalized the latency to feed (*G x Tr: F_2,43_=44.5, p<0.001*). Results are shown as scatter plots and mean ± SEM. Solid stars: comparison with the vehicle-treated *Fmr1^+/+^* group; open stars: genotype x treatment x stimulus interaction, mouse versus toy comparison; asterisks: treatment effect. One symbol: p<0.05; two symbols: p<0.01; three symbols: p<0.001. Letters: comparison with vehicle-treated *Fmr1^-/-^*group; (c): p<0.05, (b): p<0.01, (a): p<0.001. More behavioral parameters are available in Figure S6. Abbreviations: 3-Ch: three-chamber test; AAR: alternate arm returns; G: genotype effect; M: mouse; MB: marble burying test; MS: motor stereotypies; NSF: novelty-suppressed feeding test; S: stimulus effect; SAR: same arm returns; SI: social interaction test; SPA: spontaneous alternation; T: toy; Tr: treatment effect; Y-M: Y-maze test.

**Figure 4.**
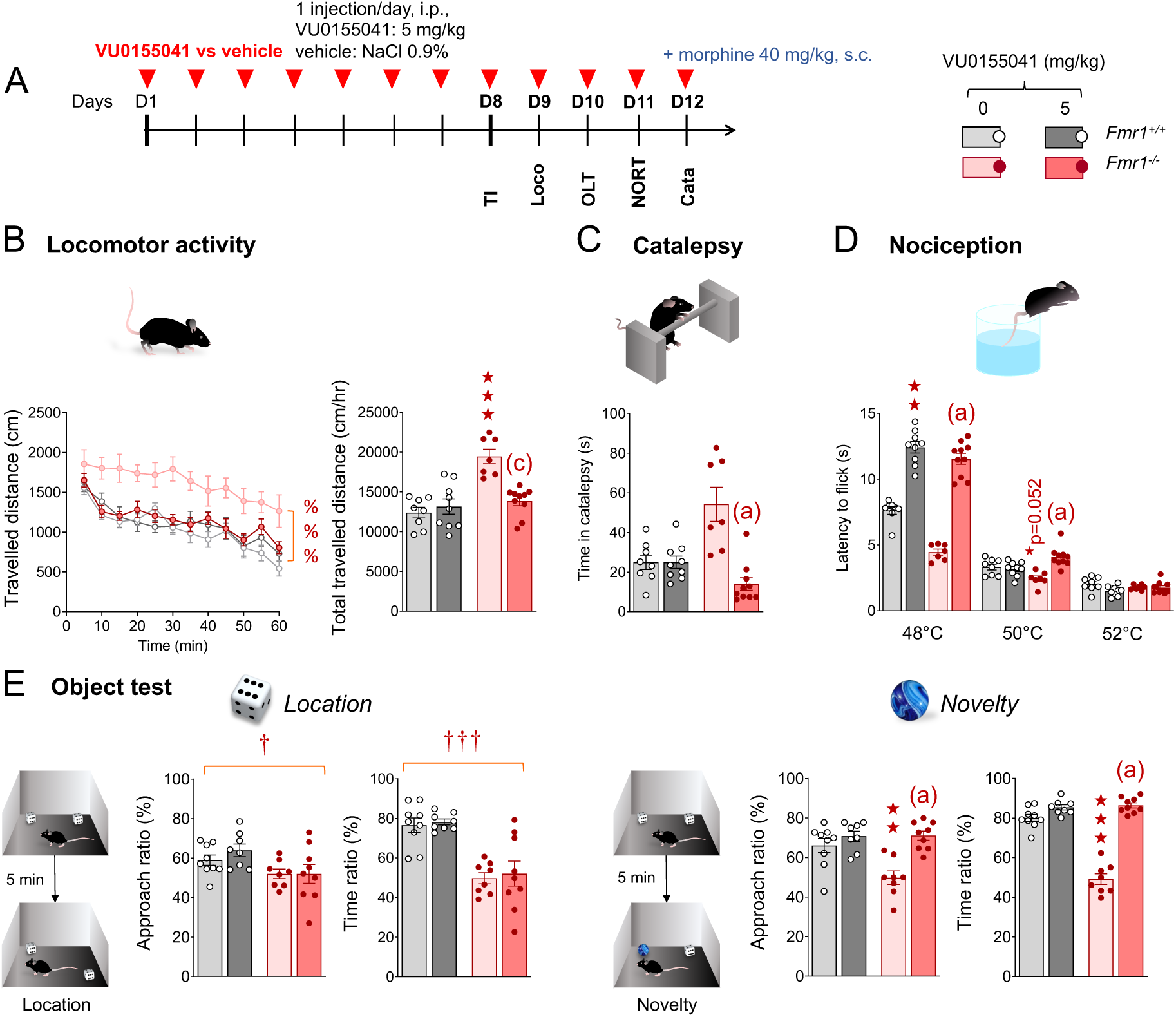
Effects of chronic facilitation of mGlu4 activity on locomotor activity, catalepsy, nociception and cognition in *Fmr1^-/-^* mice. (A) A cohort of *Fmr1^+/+^* and *Fmr1^-/-^* mice (n=8-9 per group) was injected once daily with either VU0155041 (5 mg/kg, i.p.) or vehicle (NaCl 0.9%) for 12 days. **(B)** In the locomotor activity recording test, *Fmr1^-/-^*mice displayed hyperactivity that was suppressed by VU0155041 treatment (time course: *G x Tr: F_11,330_=16.6, p<0.001,* travelled distance in 60 min: *H_3,34_=16.8, p<0.001*). (C) Following morphine administration (40 mg/kg, s.c.), *Fmr1^-/-^*mice spent longer time in catalepsy, normalized by VU0155041 (*H_3,34_=18.0, p<0.001*). (D) In the tail immersion test, VU0155041 increased nociceptive thresholds in both mouse lines at 48°C (*H_3,34_=27.3, p<0.001*). At 50°C, VU0155041 increased the latency to flick in *Fmr1^-/-^* mice only (*G x Tr: F_1,30_=16.7, p<0.001*); a tendency for decreased thresholds in *Fmr1^-/-^* mice did not reach significance (*p=0.052*). **(E)** In the object location test, *Fmr1^-/-^* mice displayed reduced approach (*G: F_1,30_=7.5, p<0.05*) and time (*G: F_1,30_=40.6, p<0.001*) ratios, without VU0155041 effects. In the novel object recognition test, the deficit detected in *Fmr1^-/-^* mice was rescued by VU0155041 administration (approach: *G x Tr: F_1,30_=7.5, p<0.05;* time*: G x Tr: F_1,30_=78.2, p<0.001*). Results are shown as scatter plots and mean ± SEM. Percents: genotype x treatment interaction; solid stars: comparison with the vehicle-treated *Fmr1^+/+^* group; daggers: genotype effect. One symbol: p<0.05; two symbols: p<0.01; three symbols: p<0.001. Letters: comparison with vehicle-treated *Fmr1^-/-^* group; (c): p<0.05, (a): p<0.001. Abbreviations: Cata: catalepsy; G: genotype effect; Loco: locomotor activity; OLT: object location test; NORT: novel object recognition test; Tr: treatment effect; TI: tail immersion.

**Figure 5.**
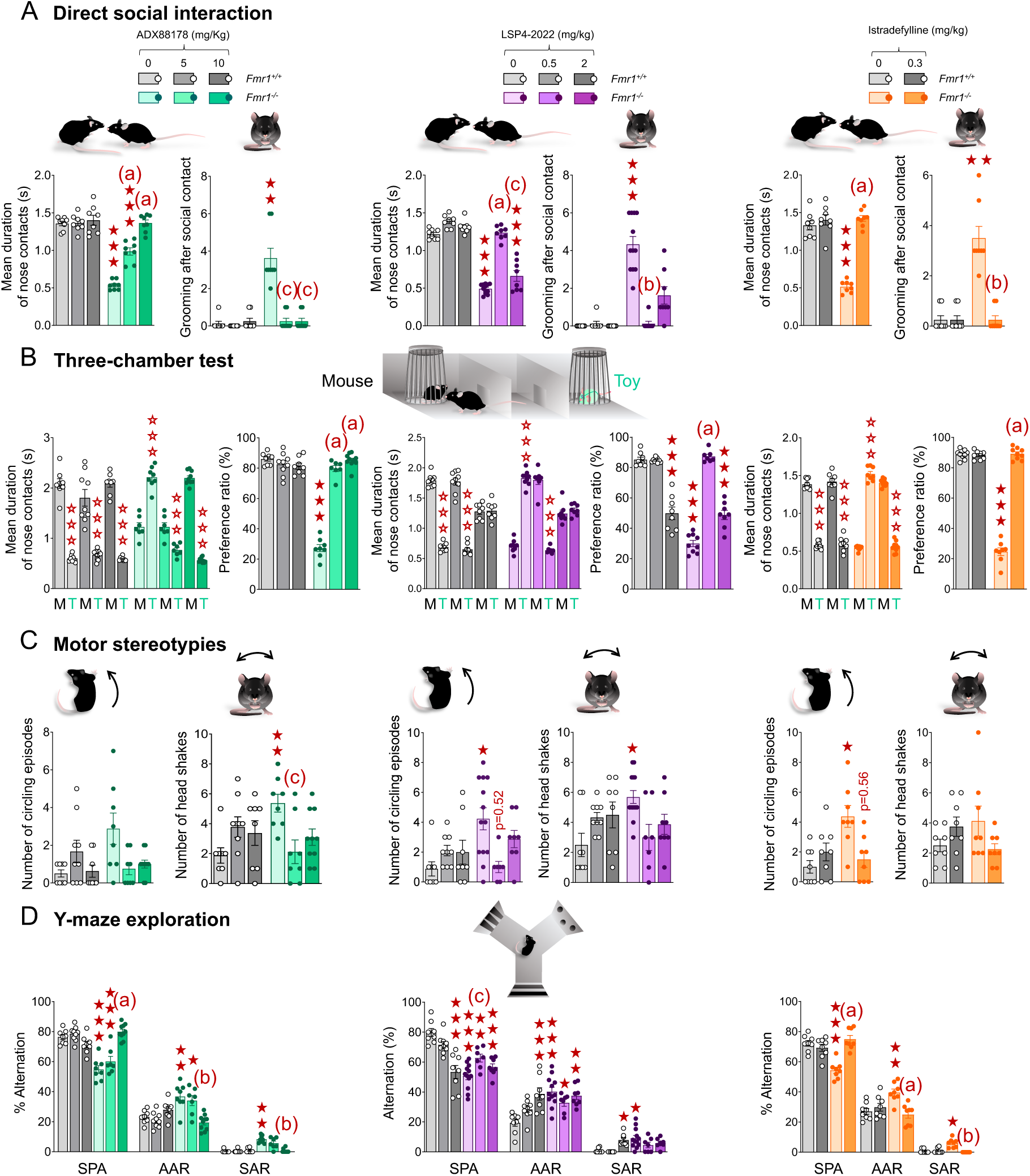
Compared effects of chronic mGlu4 facilitation or orthosteric activation, and adenosine A2A receptor blockade on core ASD-like symptoms in *Fmr1^-/-^*mice. (A) In the direct social interaction test, ADX88178 (n=8-10 per group), dose-dependently restored the mean duration of nose contacts (*G x Tr: F_2,42_=40.5, p<0.001*) and suppressed grooming after social contact since the dose of 5 mg/kg (*H_5,48_=30.4, p<0.001*). LSP4-2022 (n=8-13 per group) demonstrated beneficial effects on social parameters at 0.5 but not 2 mg/kg (nose contact duration: *G x Tr: F_2,46_=36.3, p<0.001,* grooming after social contact: *H_5,52_=43.6, p<0.001*). Istradefylline (n=8-9 per group) restored social interaction (nose contact duration: *G x Tr: F_1,28_=65.7, p<0.001,* grooming after social contact: *H_3,32_=21.5, p<0.001*). **(B)** In the three-chamber test, ADX88178 rescued social preference (duration of nose contacts: *G x Tr x S: F_2,44_=297.3, p<0.001,* preference ratio: *G x Tr: F_2,44_=150.0, p<0.001*). LSP4-2022 restored social preference at 0.5 mg/kg; both *Fmr1^+/+^* and *Fmr1^-/-^* mice treated at the dose of 2 mg/kg failed to discriminate between the mouse and the toy (duration of nose contacts: *G x Tr x S: F_2,47_=64.4, p<0.001,* preference ratio: *G x Tr: F_2,47_=42.3, p<0.001*). Istradefylline normalized social preference (duration of nose contacts: *G x Tr x S: F_1,30_=614.6, p<0.001,* preference ratio: *G x Tr: F_1,30_=442.7, p<0.001*). (C) All three compounds reduced motor stereotypies in *Fmr1^-/-^* mice (ADX88178 - head shakes: *G x T: F_2,45_=7.8, p<0.01;* LSP4-2022 - circling episodes: *H_5,53_=17.1, p<0.01;* head shakes: *H_5,53_=12.8, p<0.05;* istradefylline - circling episodes: *H_3,32_=11.9, p<0.01*). (D) In the Y-maze, ADX88178 restored SPA in *Fmr1^-/-^* mice at 10 mg/kg (*G x T: F_2,44_=31.4, p<0.001*), by reducing AAR (*G x Tr: F_2,44_=16.0, p<0.001*) and SAR (*H_5,50_=34.3, p<0.001*). Low dose of LSP4-2022 partially restored SPA (*G x Tr: F2,46=17.5, p<0.001*) in *Fmr1^-/-^* mice but failed to restore AAR (*G x Tr: F2,46=8.2, p<0.001*) and SAR (*H5,52=27.8, p<0.001*). At 2 mg/kg, LSP4-2022 altered the pattern of alternation of *Fmr1^+/+^* mice. Finally, istradefylline restored SPA (*G x Tr: F_1,28_=29.7, p<0.001*) by reducing AAR (*G x Tr: F1,28=14.0, p<0.001*) and SAR (*H_3,32_=20.8, p<0.001*) in *Fmr1^-/-^* mice. Results are shown as scatter plots and mean ± SEM. Solid stars: comparison with the vehicle-treated *Fmr1^+/+^* group; open stars: genotype x treatment x stimulus interaction, mouse versus toy comparison. One symbol: p<0.05; two symbols: p<0.01; three symbols: p<0.001. Letters: comparison with vehicle-treated *Fmr1^-/-^* group; (c): p<0.05, (b): p<0.01; (a): p<0.001. Abbreviations: AAR: alternate arm returns; D: day; G: genotype effect; M: mouse; S: stimulus effect; SAR: same arm returns; SPA: spontaneous alternation; T: toy; Tr: treatment effect. The timeline of experiments and additional behavioral parameters are available in Figure S8.

#### 2.6.1. Locomotor activity under pharmacological challenge or chronic treatment

Locomotor activity was assessed in clear Plexiglas boxes (21 × 11× 17 cm) placed over a white Plexiglas infrared-lit platform. Light intensity of the room was set at 15 lx. The trajectories of the mice were analyzed and recorded via an automated tracking system equipped with an infrared-sensitive camera (Videotrack; View Point, Lyon, France). To focus on forward activity, only movements which speed was over 6 cm/s were considered for the measurement of locomotor activity.

When monitoring the effects of dopamine receptor agonists, mice were placed in the activity boxes for a habituation period of 60 min before SKF81297 administration (to obtain stable low activity) and 15 min before quinpirole injection (to avoid a floor effect when measuring locomotor inhibition). Mice were then injected with either vehicle (NaCl 0.9%), SKF81297 (0.5, 1 or 2 mg/kg, s.c.) or quinpirole (0.05 or 0.1 mg/kg, s.c.), and locomotor activity was monitored for further 80 min (SKF81297) or 60 min (quinpirole). For testing VU0155041 effects, locomotor activity was measured over 60 min, starting 30 min after daily administration (D9) of the mGlu4 PAM treatment (5 mg/kg, i.p. versus vehicle – NaCl 0.9%). The total traveled distance was calculated for the entire time course; for quinpirole experiment, a second calculation (first 30 min) was performed to avoid counting the potential biphasic effects of low-to-moderate used doses [57].

#### 2.6.2. Catalepsy

This behavior was assessed as an indicator of striatal function [58,59]. Each animal was injected with the test compound or saline (i.p. or s.c.), and the catalepsy procedure was measured using the bar test, 30 min after haloperidol (1 mg/kg), SCH23390 (0.3 mg/kg) or sulpiride (50 mg/kg) injection, and 60 min after morphine injection (40 mg/kg). The forelimbs of each experimental mouse were placed on a cylindrical plastic bar (straw, diameter: 0.4 cm) clipped in Lego bricks 3.5 cm above the table. The time during which both forelimbs remained on the bar was recorded. The test was repeated four times (inter-trial interval: 30 s).

#### 2.6.3. Social abilities

##### Direct social interaction test

The experimental protocol was adapted from [60,61]. On testing day, a pair of unfamiliar mice (not cage mates, age-, sex-, genotype- and treatment-matched) was introduced in one of 4 square arenas (50 x 50 cm, separated by 35 cm-high opaque grey Plexiglas walls) over a white infrared floor (View Point, Lyon, France) for 10 min (15 lx). Each arena received a black plastic floor (transparent to infrared) to minimize anxiety levels. The total amount of time spent in nose contact (nose-to-nose, nose-to-flank and nose-to-anogenital region), the number of these contacts, the time spent in paw contact and the number of these contacts, grooming episodes (self-grooming), notably ones occurring immediately (<5 s) after a social contact, as well as the number of following episodes were scored *a posteriori* on video recordings (infrared light-sensitive video camera) using an ethological keyboard (Labwatcher®, View Point, Lyon, France) by trained experimenters, and individually for each animal. The mean duration of nose and paw contacts was calculated as the number of events divided by the total time spent in these events.

##### Three-chamber social preference test

The experimental protocol was adapted from [60,62]. The test apparatus consisted of a grey external Plexiglas box with transparent partitions dividing the box into three equal chambers (40 x 20 x 22.5 cm). Two sliding doors (8 x 5 cm) allowed transitions between chambers. Cylindrical wire cages (18 x 9 cm, 0.5 cm diameter-rods spaced 1 cm apart) were used to contain the mouse interactors and object (soft toy mouse) placed in the two outward chambers of the 3-chamber social preference test. The test was performed under low-light conditions (15 lx) to reduce anxiety. Stimulus *Fmr1^+/+^* mice were habituated to wire cages for 2 days before the test (20 min/day). On testing day, the experimental mouse was introduced to the middle chamber and allowed to explore the whole apparatus for a 10-min habituation phase (wire cages empty). For the social preference phase, the experimental mouse was confined back in the middle-chamber while the experimenter introduced an unfamiliar *Fmr1^+/+^* age and sex-matched mouse (8-14-week-old, grouped housed) into a wire cage in one of the side-chambers and a soft toy mouse (8 x 10 cm) in the second wire cage. Then, the experimental mouse was allowed to explore the apparatus for 10 min. The sliding doors were reopened allowing an additional 10-min exploration. The time spent in each chamber and in nose contact with each wire cage, the number of these contacts and the number of entries in each chamber were scored *a posteriori* on video recordings using an ethological keyboard (Labwatcher®, View Point, Lyon, France) by trained experimenters. The mean duration of nose contacts (nose to nose, nose to flank, nose to anogenital region) was calculated from these data. Preference ratio was calculated as follows: Time in nose contact with the mouse / (Time in nose contact with the mouse + Time in nose contact with the object) x 100. The relative position of stimulus mice was counterbalanced between groups.

#### 2.6.4. Stereotyped and perseverative behaviors

##### Motor stereotypies

The experimental protocol was adapted from [63]. To detect spontaneous motor stereotypies in *Fmr1^-/-^* versus *Fmr1^+/+^*animals, mice were individually placed in clear standard home cages (21 × 11 × 17 cm) filled with 3-cm deep fresh sawdust for 10 min. No water was available. Light intensity was set at 30 lx. Trained experimenters scored numbers of spontaneous head shakes, rearing, burying, grooming and circling episodes and the total amount of time spent burying by direct observation.

##### Marble-burying

Marble burying was used as a measure of stereotyped/perseverative behavior [62,64]. Mice were introduced individually in transparent cages (21 × 11 × 17 cm) containing 20 glass marbles (diameter: 1.5 cm) evenly spaced on 4-cm deep fresh sawdust. To prevent escapes, each cage was covered with a filtering lid. Light intensity in the room was set at 40 lx. The animals were removed from the cages after 15 min, and the number of marbles buried more than half in sawdust was recorded.

##### Y-maze exploration

Spontaneous alternation behavior was used to assess perseverative behavior [65,66]. Each Y-maze (Imetronic, Pessac, France) consisted of three connected Plexiglas arms (15 x 15 x 17 cm) covered with distinct wall patterns (15 lx). Floors were covered with lightly sprayed fresh sawdust to limit anxiety. Each mouse was placed at the center of the maze and allowed to freely explore this environment for 5 min. The pattern of entries into each arm was quoted on video-recordings. Spontaneous alternations (SPA), i.e. successive entries into each arm forming overlapping triplet sets, alternate arm returns (AAR) and same arm returns (SAR) were scored, and the percentage of SPA, AAR and SAR was calculated as following: total / (total arm entries -2) * 100.

#### 2.6.5. Anxiety-like behavior

##### Novelty-suppressed feeding

The protocol was adapted from [67]. Novelty-suppressed feeding (NSF) was measured in 24-hr food-deprived mice, isolated in a standard housing cage for 30 min before individual testing. This test was performed in the same arenas as the ones used for direct social interaction. Three pellets of ordinary lab chow were placed on a white tissue in the center of each arena, lit at 60 lx. Each mouse was placed in a corner of an arena and allowed to explore for a maximum of 15 min. Latency to feed was measured as the time necessary to bite a food pellet. Immediately after an eating event, the mouse was transferred back to home cage (free from cage-mates) and allowed to feed on lab chow for 5 min. Food consumption in the home cage was measured.

#### 2.6.6. Nociceptive thresholds

##### Tail-immersion test

This test was performed as previously described [41,68]. Nociceptive thresholds were assessed by immersing the tail of the mice (5 cm from the tip) successively into water baths at 48°C, 50°C and 52°C. Mice were gently maintained in a tissue pocket during this experiment. The latency to withdraw the tail was measured at each temperature, with a cut-off of 10 s.

#### 2.6.7. Cognitive performance

##### Object location and novel object recognition test

The experimental paradigm was adapted from [69]. The test was performed on two successive days. On day 1, during familiarization phase (10 min), the mice were presented with two copies of an unfamiliar object. The animals were returned to their home cage for a 5 min intertrial interval. During this interval, the experimenter displaced one of the two objects to a novel location in the arena, before the mice were allowed to freely explore the apparatus for 10 min (location phase). On day 2, during familiarization phase (10 min), the mice were presented a pair of novel unfamiliar objects. They were returned to their home cage for 5 min. During this interval, the experimenter replaced one of the two objects by a novel object. Then the mice were allowed to explore freely the arena for another 10 min (novelty phase). Stimuli objects sized 1.5-3 x 2-3 cm. The identity of the objects and their spatial location were balanced between subjects. The number of visits and the time spent exploring each object were scored *a posteriori* on video recordings. A percentage of discrimination was calculated for number of visits and time exploring the objects as following: exploration of displaced or novel object / total exploration * 100.

### 2.7. Photopharmacology experiments

#### 2.7.1. Surgery

Both 2-months-old male and female mice were used in this experiment. After surgery, mice were singly housed under a 12h light-dark cycle with food and water ad libitum. Surgeries were performed under chemical anesthesia (ketamine at 50 mg/kg and xylazine at 7.5 mg/kg). We used an optofluidic device (OsFC_240/250-0.63_5mm_FLT_100/170_0.5; Doric Lenses, Quebec, QC, Canada) to both deliver the optogluram, a light-sensitive mGlu4 PAM, and control it by light. The injection cannula (inner diameter: 100 μm; outer diameter 170 μm) coupled to an optic fiber (inner diameter: 240 μm; outer diameter 250 μm, numerical aperture = 0.63) was implanted unilaterally in the ventral pallidum at the following stereotaxic coordinates: +0.55 mm (AP); +/-1.3 mm (ML: counterbalanced right and left hemispheres); -4.9 mm (DV). It was fixed on the skull by using dental acrylic cement. Mice were returned to their home cages and placed under a heating light for 24h post-surgery and i.p. ketoprofen injections (2 mg/kg) were administrated for analgesia until the end of recovery. Mice with uncorrected targeting of fibers were excluded from this experiment (n=2, one *Fmr1^-/-^*and one *Fmr1^+/+^*).

#### 2.7.2. Photopharmacology

We used the photopharmacology tool to modulate specifically postsynaptic mGlu4 located on NAc D2-SPNs terminals in the VP combining two wavelengths coupled to a rotating optical fiber. Each channel was controlled via the LED driver software (Doric Lenses, Quebec, QC, Canada). After 3 weeks of recovery, mice were habituated to the optic fiber connection one week before the tests.

Optogluram (30 µM), a light-sensitive mGlu4 PAM, was injected unilaterally in the VP, through the cannula coupled with optic fiber 20 min before starting the behavior test. We controlled optogluram activity by switching it from active (resting) to inactive state (by delivering violet light, 380 nm wavelength, 50 ms light pulses at 10 Hz) state, then back to active state (green light, 500 nm wavelength, 50 ms light pulses at 10 Hz). Social behavior was assessed in *Fmr1^+/+^*and *Fmr1^-/-^* mice using the direct social interaction test. Fiber implanted mice were introduced with an unfamiliar mouse (sex, age, genotype-matched, not implanted) in a square arena (50 x 50 cm, separated by 35 cm-high opaque grey Plexiglas walls) over a white infrared floor (View Point, Lyon, France) for 9 min (15 lx). The test was divided in three phases: the 3-min baseline, when the light was OFF and optogluram active, the following 3-min, when the violet light was ON and optogluram inactive and the last 3-min when the light was switched to green and optogluram returned to its active state. After another 3 weeks, all the mice underwent the same protocol after intra-VP injection of vehicle, to assess the effects of light *per se*. The same parameters as in the direct social interaction test were scored. Finally, brains were collected to verify cannula implantation sites.

### 2.8. Statistics

Statistical analyses were performed using Statistica 9.0 software (StatSoft, Maisons-Alfort, France) or Prism 10.6 (GraphPad Software, USA). For all comparisons, values of p<0.05 were considered as significant. If normality of residuals was respected, statistical significance in behavioral experiments was assessed using two-way analysis of variance (followed by Tukey’s post-hoc test in case of significant genotype x treatment interaction). In case of non-normality, we used the non-parametric Mann-Whitney U test or Kruskal-Wallis analysis of variance (followed by 2-tailed multiple comparison of mean ranks); consequently, genotype and treatment effects could not be interpreted in such two-factors experimental designs. In case of repeated or matched measures, a repeated three-way analysis of variance (time course of locomotor activity, three-chamber test) or a linear mixed-effects model (photopharmacology, RNAscope® and qRT-PCR experiments) was performed (followed by Tukey’s post-hoc test in case of significant interaction). Unsupervised clustering analysis was performed on transformed qRT-PCR data [30,70] using complete linkage with correlation distance (Pearson correlation) for drug, treatment and brain region (Cluster 3.0 and Treeview software). Based on *a posteriori* multiple logistic regression (Table S2), principal component analysis (Figure S1) and principal component logistic regression (Table S3), both male and female groups showed similar behavior. Therefore, data of male and female subjects were pooled in the present study.

## 3. Results

### 3.1. Altered striatal-dependent behaviors in Fmr1^-/-^ mice

To test the hypothesis of altered striatal function in *Fmr1^-/-^*mice, we evaluated striatal-dependent behaviors. First, we assessed the stimulant locomotor effects of the dopamine D1/D5 dopamine receptor agonist SKF81297 (s.c.; 0, 0.5, 1 and 2 mg/kg) and the inhibitory locomotor effects of low-to-moderate doses of the D2/D3 dopamine receptor agonist quinpirole (s.c.; 0, 0.05 and 0.1 mg/kg) in *Fmr1^-/-^*and *Fmr1^+/+^* mice (Figure 1A). Consistent with previous reports [71,72], locomotor activity during habituation in both experiments was higher in *Fmr1^-/-^* mice as compared to *Fmr1^+/+^*mice. Under SKF81297 administration, *Fmr1^-/-^* mice displayed increased locomotor response compared to *Fmr1^+/+^* mice, regardless of the dose. Both doses of quinpirole inhibited locomotion in *Fmr1^+/+^*mice for the first 30 minutes. However, this effect was maintained up to 60 minutes for the low dose (0.05 mg/kg) only (Figure S2), while the dose of 0.1 mg/kg may have shown a biphasic effect [57]. In contrast, quinpirole-induced suppression of locomotor activity was impaired in *Fmr1^-/-^* mice. Together, these results indicate that D1/D5 receptor agonist-induced activation is increased, whereas D2/D3 receptor activation is weaker in the absence of FMRP.

We then evaluated the cataleptic response induced by dopamine receptor antagonists, either targeting D1/D5 and D2/D3 receptors indifferently (haloperidol, 1 mg/kg), or targeting preferentially D1/D5 (SCH23390, 0.3 mg/kg) or D2/D3 (sulpiride, 50 mg/kg) (Figure 1B). Regardless of the molecule, *Fmr1^-/-^* mice spent less time in an unnatural position than *Fmr1^+/+^* mice. In contrast, when injected with a high dose of the opioid receptor agonist morphine (40 mg/kg), *Fmr1^-/-^* mice spent more time under cataleptic state than *Fmr1^+/+^* mice. Taken together, these results suggest a differential sensitivity of the striatum to dopamine receptor activation in *Fmr1^-/-^*versus *Fmr1^+/+^* mice.

### 3.2. Deficient social interaction and decreased number of Fos-positive D1-SPNs in the striatum of Fmr1^-/-^ mice

Altered striatal-dependent behavior in *Fmr1^-/-^* mice could result from excessive D2-SPN output activity, a potential neurobiological substrate for impaired social behavior in this ASD model [30]. Here, we used RNAscope ISH to assess the activity (*Fos* labelling) of D1-(*Pdyn* labelling) and D2-(*Penk* labelling) SPNs in the DS (medial and lateral) and NAc (Core and Shell) in *Fmr1^-/-^* and *Fmr1^+/+^* mice 45 min after a social interaction test (Figure 2A).

Consistent with previous findings, *Fmr1^-/-^* mice displayed a severe deficit in social interaction (Figure 2B, S3A). RNAscope ISH revealed that the percentage of *Pdyn^+^-Fos*^+^ double-labelled cells (activated D1-SPNs) was reduced in the whole striatum of *Fmr1^-/-^*mice compared to *Fmr1^+/+^* mice. Meanwhile, the proportion of *Penk^+^-Fos*^+^ double-labelled cells (activated D2-SPNs) was found unchanged across striatal regions. Accordingly, the ratio of *Pdyn^+^-Fos*^+^ over *Penk^+^*-*Fos*^+^ labelled striatal cells was decreased in *Fmr1^-/-^* mice (Figure 2C-D). Although the expression of both *Pdyn* and *Penk* was decreased in *Fmr1^-/-^*mice (Figure S4), the total number of *Pdyn^+^*or *Penk^+^* cells was similar (Figure S3B) indicating that neither the expression of SPN markers nor the total number of cells influenced the ratio of double-labelled cells. Altogether, these results suggest that the D1/D2-SPN balance of activity in *Fmr1^-/-^*mice was shifted towards deficient striatal D1-SPN activity and, therefore, that D2-SPN outputs outweighed D1-SPN outputs following social interaction in these mice.

### 3.3. Rescued social behavior in Fmr1^-/-^ mice under chronic facilitation of mGlu4 activity

Considering D2-SPN activity is likely to be excessive in the striatum of *Fmr1^-/-^* mice, we reasoned that treatment with a pharmacological inhibitor of D2-SPN activity such as a mGlu4 PAM would reverse their behavioral deficits. We first verified *in cellulo* that the absence of *Fmr1* in HEK293 cells does not affect the expression of G protein-coupled receptors and their coupling to G proteins. In *Fmr1^-/-^* cells, mGlu4 show similar levels of expression and function as in *Fmr1^+/+^* HEK293 cells (Figure S5). In *Fmr1^-/-^*mice, *Grm4* mRNA expression was maintained compared to *Fmr1^+/+^* mice across several brain regions (Figure S4). We then evaluated the effects of chronically administering (D1 to D18) the mGlu4 PAM VU0155041 (0, 2.5 or 5 mg/kg) on ASD-like behavioral deficits in *Fmr1^-/-^* and *Fmr1^+/+^* mice (Figure 3A).

We first focused on social behavior. In the direct social interaction test (Figure 3B and Figure S6A), *Fmr1^-/-^* mice displayed a severe deficit that was reversed under chronic VU0155041 treatment, partially at the dose of 2.5 mg/kg, and fully at the dose of 5 mg/kg. When VU0155041 was administered only once, this 5 mg/kg dose produced only partial effects on social interaction (Figure S7). In the 3-chamber test for social preference (Figure 3C and Figure S6B), *Fmr1^-/-^* mice failed to display preference for spending more time interacting with a congener over a toy. VU0155041 restored social preference from the lowest dose tested in *Fmr1^-/-^* mice. Together, these data indicate that facilitating mGlu4 activity alleviates social deficits in *Fmr1^-/-^*mice.

### 3.4. Normalized stereotypic behavior, anxiety, activity, catalepsy and cognition in Fmr1^-^/- mice under chronic facilitation of mGlu4 activity normalized

We then assessed the effects of chronic VU0155041 on non-social ASD-related stereotypic motor behavior, perseveration and anxiety-like behavior in *Fmr1^-/-^* and *Fmr1^+/+^* mice [70–72].

Regarding spontaneous motor stereotypies, VU0155041 at both tested doses suppressed excessive grooming and circling in *Fmr1^-/-^*mice (Figure 3D and S6C). In the marble burying test, chronic VU0155041 administration abolished excessive marble burying in *Fmr1^-/-^* mice from the lowest dose (2.5 mg/kg) (Figure S6D). During Y-maze exploration, *Fmr1^-/-^* mice displayed reduced spontaneous alternation (SPA) rate and increased perseverative same arm re-entries (SAR) compared to *Fmr1^+/+^* mice. VU0155041 suppressed SAR and normalized SPA (at the highest dose) (Figure 3E and S6E). Finally, *Fmr1^-/-^* mice displayed longer latencies to eat and reduced food intake in the novelty-suppressed feeding test, suggesting increased levels of anxiety.

Maximal efficiency of chronic VU0155041 in relieving this excessive anxiety was reached from the lowest dose (Figure 3F and S6F).

We then evaluated whether the beneficial effects of the optimal 5 mg/kg dose of VU0155041 administered chronically would generalize to other behavioral features in *Fmr1^-/-^* mice: locomotor hyperactivity, facilitated morphine-induced catalepsy, altered nociceptive thresholds and cognitive impairments [73,74] (Figure 4A).

Regarding locomotion, we confirmed hyperactivity in vehicle-treated *Fmr1^-/-^* mice over 60 min. VU0155041 administration normalized activity in these mice to *Fmr1^+/+^* levels (Figure 4B). As previously observed (Figure 1B), vehicle-treated *Fmr1^-/-^* mice exhibited longer catalepsy following morphine administration than *Fmr1^+/+^* mice; mGlu4 facilitation suppressed this difference (Figure 4C). In the tail immersion test, VU0155041 increased nociceptive thresholds in both *Fmr1^-/-^* and *Fmr1^+/+^* mice, at 48°C. At 50°C, analgesia was only observed in VU0155041-treated compared to vehicle-treated *Fmr1^-/-^* mice (Figure 4D). Lastly, *Fmr1^-/-^* mice displayed a deficit both in detecting the change in location of an object and recognizing a novel object. Chronic VU0155041 failed to restore object location performance but fully rescued novel object recognition in these mice (Figure 4E). These results indicate that the benefit of mGlu4 PAM administration exceed ASD-like symptoms in *Fmr1^-/-^* mice and extend to other, more FXS-specific, behavioral comorbidities.

### 3.5. Beneficial behavioral effects in Fmr1^-/-^ mice treated with different pharmacological compounds known to dampen D2-SPN activity

To further explore the interest of facilitating mGlu4 activity to relieve behavioral deficits in *Fmr1^-/-^* mice, we evaluated the effects of chronic treatment with another mGlu4 PAM, ADX88178 (0, 5 or 10 mg/kg), and the orthosteric mGlu4 agonist LSP4-2022 [51] (0, 0.5 or 2 mg/kg). Moreover, we assessed the effects of a different inhibitor of D2-SPN activity, the adenosine 2A receptor (A2A) antagonist istradefylline [46] (0.3 mg/kg) (see timeline of experiments and more behavioral parameters in Figure S8).

Chronic administration of ADX88178 demonstrated similar efficiency as VU0155041 in relieving social behavior deficits, stereotypies, and perseveration in *Fmr1^-/-^* mice. In contrast, treatment with LSP4-2022 demonstrated beneficial effects at the low dose of 0.5 mg/kg only; at a higher dose (2 mg/kg), this compound not only failed to alleviate ASD-like behavior in *Fmr1^-/-^* mice but showed deleterious effects on social and perseverative behavior in *Fmr1^+/+^*mice, possibly by activating mGlu7 receptors [75]. Last of all, the A2A antagonist istradefylline (0.3 mg/kg) improved both social and repetitive behavior in *Fmr1^-/-^* mice (Figure 5A-D). Overall, these results indicate that blocking the A2A receptor shows similar efficacy in alleviating ASD-like symptoms as facilitating mGlu4 activity with a higher dose of both PAMs, VU0155041 and ADX88178, or activating mGlu4 with a lower dose of the agonist LSP4-2022 (Figure S1).

### 3.6. Alleviated social deficits in Fmr1^-/-^ mice under optical modulation of presynaptic mGlu4 in the ventral pallidum

In previous experiments, mGlu4 PAMs/agonists were administered peripherally, targeting mGlu4 in the whole brain and organism. To better dissect the contribution of NAc D2-SPN inhibition to therapeutic effects in *Fmr1^-/-^* mice, we used photopharmacology to modulate the activity of mGlu4 expressed in the projection site of NAc D2-SPN terminals, the VP. We injected optogluram, a light-sensitive mGlu4 PAM [76], in this region, through a cannula coupled with an optic fiber. We controlled optogluram activity by switching it from active (resting) to inactive (violet light, 380 nm) state, then back to active state (green light, 500 nm) (Figure 6A, B). After 3 weeks of post-surgery recovery, social behavior was assessed in *Fmr1^+/+^* and *Fmr1^-/-^* mice using the direct social interaction test (Figure 6C). After another 3 weeks, all the mice underwent the same protocol after intra-VP injection of vehicle, to assess the effects of light *per se* (microinjection of optogluram vehicle, Figure S9C).

**Figure 6.**
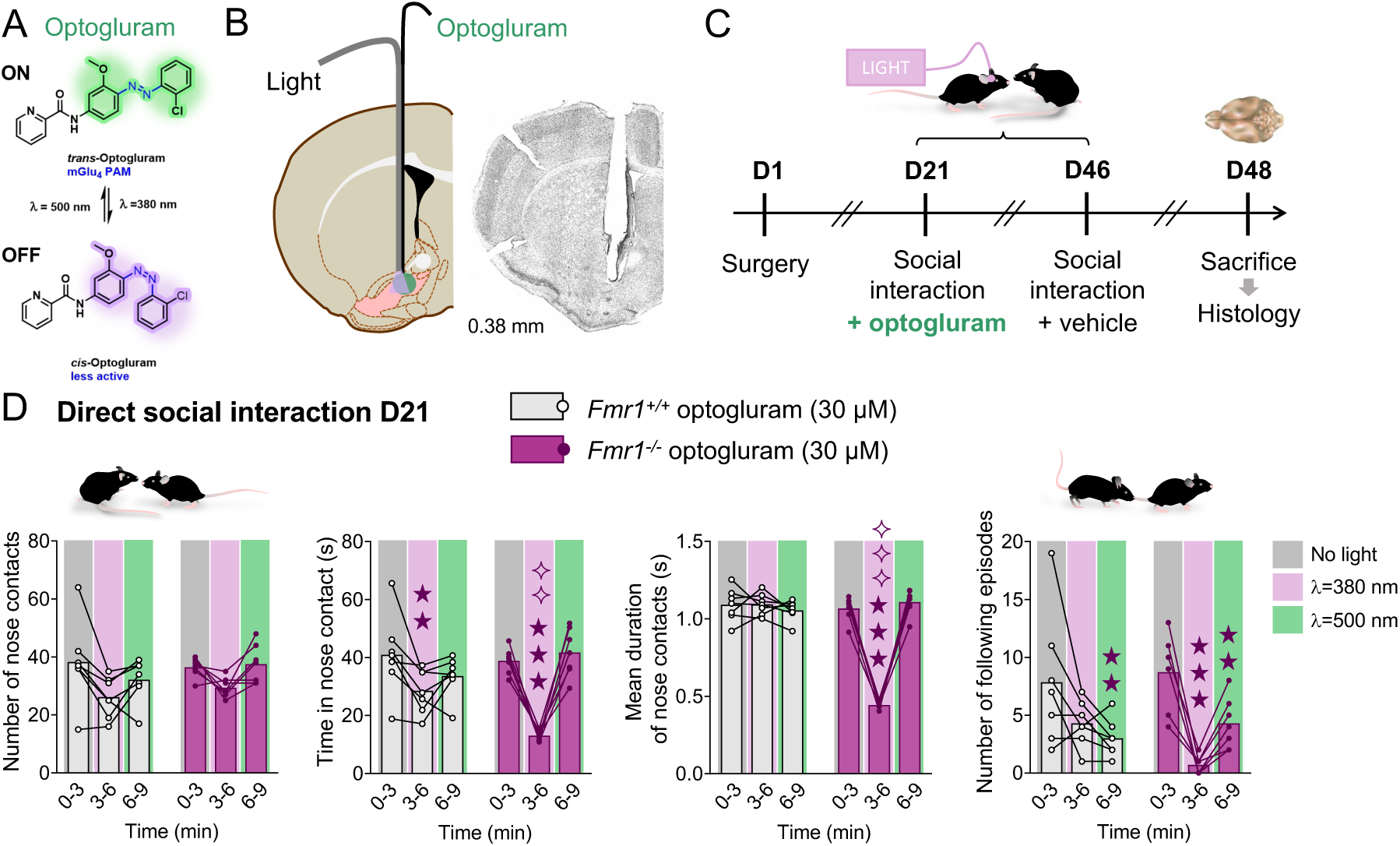
Effects of photopharmacological activation within the ventral pallidum of the mGlu4 PAM optogluram on social interaction in *Fmr1^-/-^*mice. **(A)** The mGlu4 PAM optogluram can be switched from a trans active configuration to a cis-inactive configuration after illumination with violet light (λ=380 nm) and from cis to trans with green light (λ=500 nm). **(B)** Scheme: a hybrid cannula for light and fluid delivery was positioned at the upper limit of VP. Picture: representative microphotograph showing cannula placement in the VP. Coordinates refer to bregma. Cannula placement for all mice was mapped on schematic brain sections in Figure S9D. **(C)** Timeline of the experiments. A first social interaction session was performed in presence of optogluram at D21 following surgery (n=7 mice per group). A second session was performed at D46 during which the vehicle of optogluram (NaCl 0.9%) was microinjected into the VP (Figure S9C). Mice were sacrificed at D48 for verification of cannula placement. **(D)** In the direct social interaction test, under no light condition (0-3 min), optogluram was active and *Fmr1^-/-^* mice behaved similarly as *Fmr1^+/+^* mice. Illuminating optogluram with violet light (3-6 min) reduced the number of nose contacts regardless of the treatment and genotype (*A: F_2,24_=10.9, p<0.001*). This manipulation inactivated optogluram and reduced the time spent in nose contact in both *Fmr1^+/+^* and *Fmr1^-/-^* mice, but more severely in the latter. Green light, by reactivating optogluram (6-9 min), normalized the time of interaction in *Fmr1^-/-^* mice back to *Fmr1^+/+^*mice levels (*G x Tr x A: F_2,24_=12.0, p<0.001*). Inactivating optogluram markedly reduced the mean duration of nose contacts in *Fmr1^-/-^* and not *Fmr1^+/+^* mice, and reactivating optogluram rescued this parameter (*G x Tr x A: F_2,24_=51.0, p<0.001*). Switching off optogluram decreased the number of following episodes in *Fmr1^-/-^* mice (*G x Tr x A: F_2,24_=4.4, p<0.05*); reactivation under green light tended to restore this behavior (p=0.062) while a decrease was observed in *Fmr1^+/+^* animals. Results are shown as scatter plots and mean ± SEM. Solid stars: comparison with the optogluram-treated *Fmr1^+/+^* group under no light conditions; diamonds: comparison with the optogluram-treated *Fmr1^+/+^* group under the same light conditions. One symbol: p<0.05; two symbols: p<0.01; three symbols: p<0.001. Abbreviations: A: optogluram activation (on versus off); D: day; G: genotype; Tr: treatment (optogluram versus vehicle; vehicle data are displayed in Figure S9C).

During the 3-min baseline, when the light was off and optogluram active, *Fmr1^-/-^*mice displayed normalized social behavior. When violet light (380 nm) was switched on and optogluram inactivated (3-6 min), a significant deficit in social interaction became detectable in *Fmr1^-/-^*mice, which was completely rescued under green light (500 nm) and reactivation of optogluram (6-9 min) (Figures 6D and S9B). Switching on and off optogluram reduced the time spent in nose contact in *Fmr1^+/+^*mice, without effect on mean duration of nose contacts. Moreover, neither green or violet light in the absence of optogluram influenced behavior in *Fmr1^+/+^* and *Fmr1^-/-^* mice (Figure S9C). Thus, facilitating the activity of presynaptic mGlu4 in the VP restored social abilities in *Fmr1^-/-^* mice.

## 4. Discussion

FXS is currently the leading monogenic cause of ASD [4]. The striatum plays a key role in the ASD-like phenotype [23,24]. To date, clinical [77,78] and preclinical studies [79–82] investigating striatal dysfunction in FXS have mainly focused on the DS and its cortical afferences. However, social withdrawal, reward deficit, repetitive behaviors, and lack of flexibility in FXS and its *Fmr1^-/-^*mouse model [4,21] point towards ventral striatal dysfunction [22,28,30–34]. Consistent with this, altered physiology and connectivity of D1- and D2-SPNs have been evidenced in the NAc of *Fmr1^-/-^* mice [37–40]. In the present study, we interrogated a potential imbalance of D1/D2-SPN activity in *Fmr1^-/-^* mice, notably in the NAc, and its potential therapeutic implications for the management of ASD symptomatology.

We first assessed the locomotor effects of dopaminergic agonists in *Fmr1^-/-^* mice, as a proxy of striatal function. The D1/D5 receptor agonist SKF81297 dose-dependently produced hyperlocomotion in mice, and this hyperlocomotion was more pronounced in *Fmr1^-/-^* mice, consistent with increased mEPSC and synaptic contacts at dorsolateral D1-SPNs in *Fmr1^-/-^* mice [82]. In contrast, low doses of the D2/D3 receptor agonist quinpirole inhibited locomotion less in *Fmr1^-/-^* mice than in their *Fmr1^+/+^* controls. At low doses, quinpirole was reported to reduce locomotion in mice [57,83,84]. This inhibition would mainly result from the activation of D2 presynaptic and/or somatodendritic autoreceptors at mesencephalic DA neurons and the subsequent decrease in striatal DA release [84–86]. In *Fmr1^-/-^* mice, although *Drd2* expression was not regulated in either the VTA or striatum (Figure S4), an increase in striatal D2 receptor density was reported [87]. Besides, increasing D2 receptor expression in NAc Core D2-SPNs is sufficient to increase ambulatory behavior in mice [88]. Thus, an increase of somatodendritic D2 receptor density in the NAc of *Fmr1^-/-^*mice may have reduced the efficacy of low doses of D2 receptor agonist. We then explored catalepsy in *Fmr1^-/-^*mice. The cataleptic response to D1/D5 and D2/D3 receptor antagonists was reduced in *Fmr1^-/-^* mice, possibly a consequence of locomotor hyperactivity in *Fmr1^-/-^* mice [71,72]. Conversely, akinetic and cataleptic effects of morphine were increased in *Fmr1^-/-^* mice, indicating an increased morphine-induced response of the NAc, where the *Oprm1* gene, coding for mu opioid receptor, is enriched and expressed in both types of SPNs [89–91]. A2A receptor antagonists reduce morphine-induced catalepsy [92,93], likely by reducing NAc D2-SPN activity [45], which indirectly points to increased NAc D2-SPN activity in *Fmr1^-/-^* mice.

We then used RNAscope® to assess striatal D1/D2-SPN activity balance in *Fmr1^-/-^* mice, following a social encounter. We observed that the proportion of *Pdyn*-*Fos-* positive neurons, namely active D1-SPNs, was lower in the striatum, DS and NAc, of *Fmr1^-/-^* mice compared to *Fmr1^+/+^* mice, resulting in decreased *Pdyn*/*Penk*-*Fos*-positive ratio. This modification of the ratio was not due to neuronal loss or change in the expression of SPN markers, since the total number of *Pdyn*-positive SPNs was maintained and both *Pdyn* and *Penk* expression were decreased in *Fmr1^-/-^* mice. Whether this decrease would be less pronounced in the DS compared to NAc will need further investigation, with more statistical power. Our finding of decreased *Fos*-labelled D1-SPNs, notably in the NAc, matches recent report of decreased D1-SPN excitability, but increased D2-SPN excitability, in the NAc Core of *Fmr1^-/-^* mice [39]. One might propose that such changes would result from compensatory mechanisms in response to protein expression dysregulations (e.g. dopamine receptors) in the absence of FMRP [82,87]. Either way, modifications in SPN excitability likely involves modified synaptic transmission within cortico- and amygdalo-striatal circuits, and notably increased synaptic strength within the BLA/PFC to NAc Core pathways [40,82]. NAc D2-SPNs were shown to play a critical role in controlling social behavior. Their activation in naive mice make them more susceptible to social stress, and socially defeated mice, which display social avoidance, show increased NAc D2-SPN activity [33]. Accordingly, an overweight of NAc D2-SPN outputs, resulting either from reduced NAc D1-SPN outputs or from direct stimulation of NAc D2-SPNs, induces social avoidance, while activating NAc D1-SPNs facilitates social behavior [30,34]. Hence, we propose that an imbalance between NAc SPN activity, in favor of D2-SPNs, contributes to the social behavior deficit observed in *Fmr1^-/-^*mice.

Consistent with this hypothesis, treating *Fmr1*^-/-^ mice chronically with pharmacological compounds that can suppress D2-SPN activity relieved their ASD-like behavioral deficits. The mGlu4 PAMs VU0155041 and ADX88178 restored social interaction and social preference in a dose-dependent manner and reduced stereotypic and perseverative behaviors in *Fmr1^-/-^* mice. The orthosteric mGlu4 agonist LSP4-2022 produced similar beneficial effects, but at low dose only, when more selectively recruiting mGlu4 [94]. Thus, facilitating mGlu4 signaling rescues ASD-like behavior in *Fmr1^-/-^* mice, as previously shown in this model [47], in other mouse models of ASD [95,96] or in mice with excessive NAc D2-SPN activity [30,41]. Restoration of social behavior is expected to involve inhibition of NAc D2-SPNs, while reduced stereotypies may result from inhibition of D2-SPN populations in the dorsomedial, central and ventral striatum (i.e. NAc) [30,97,98]. However, the imbalance between SPN activity appears to be reversed in the dorsolateral striatum of *Fmr1^-/-^* mice, in which D1-SPN excitability was reported to be increased and correlated with repetitive behaviors [82]. A finely tuned balance SPN activity likely contributes to control motor behaviors in different subregions of the DS [99].

Besides its effects on ASD-like deficits, enhancing mGlu4 activity with VU0155041 reduced hyperactivity, anxiety-like behaviors, normalized morphine-induced catalepsy, showed analgesic effects and restored novel object recognition (but not object location) in *Fmr1^-/-^* mice. Such effects may involve a widespread action of the drug at cerebral mGlu4 after systemic administration, within corticostriatal projections [43,100], hippocampus, entorhinal cortex, thalamic nuclei [43,101] and amygdala and/or spinal cord [76,102], in addition to the cerebellum [47]. The therapeutic potential of mGlu4 facilitators thus extends beyond the diagnostic symptoms of ASD in FXS, to comorbid conditions such as hyperactivity, and cognitive deficits.

When used at a high dose (2 mg/kg), recruiting mGlu7 in addition to mGlu4, the agonist LSP4-2022 showed no effect on social interaction and only partial benefit on social preference in *Fmr1^-/-^* mice, but a deleterious impact on social behavior in *Fmr1^+/+^* mice. This result suggests that activating mGlu7 may be inefficient to relieve social deficit in *Fmr1^-/-^* mice, in contrast with positive outcomes on protein synthesis and cognitive performance [103]. However, the mGlu7 PAM VU0422288 improved social preference and cognitive abilities in the *Mecp2* heterozygous (*Mecp2^+/-^*) mouse model of Rett syndrome, which displays reduced levels of mGlu7 protein expression [104]. Intriguingly, VU0422288 also behaves as a potent mGlu4 PAM [105], suggesting that a delicate interplay between mGlu4 and mGlu7 signaling may intervene in the control of social behavior, as proposed for motor control [44]. Further investigations are required to comprehensively explore the therapeutic potential of targeting mGlu7 in mouse models of ASD.

Finally, chronically administered A2A antagonist istradefylline normalized ASD-like behaviors in *Fmr1^-/-^* mice similarly to mGlu4 facilitators/activators. Istradefylline was previously found to improve novel object recognition and hippocampal function in *Fmr1^-/-^* mice [48]. Positive effects on social behavior have likely involved istradefylline-mediated inhibition of D2-SPNs, in which A2A receptor expression is highly enriched [45,46], as for mice bearing a specific ablation of NAc D1-SPNs [30]. Our results thus converge towards the therapeutic potential of compounds reducing D2-SPN activity to relieve ASD deficits in FXS. Systemic administration of these compounds, however, was not sufficient to conclude for a specific contribution of D2-SPN inhibition, notably in the NAc, to their beneficial action on ASD-like behavior in *Fmr1^-/-^* mice.

Based on high expression of mGlu4 in the cerebellum [43,106] and improved cerebellum-dependent behavioral responses in VU0155041-treated *Fmr1^-/-^* mice [47], Martin et al. proposed that positive effects of mGlu4 facilitation on social behavior may involve cerebellar mGlu4. In the present study, however, photoactivation of the mGlu4 PAM optogluram in the VP fully rescued social interaction in *Fmr1^-/-^* mice, suggesting mGlu4 in the VP are sufficient to control social behavior. This result is consistent with our hypothesis of NAc D2-SPN inhibition as a powerful pharmacological lever to restore social behavior in the *Fmr1^-/-^* mouse model. The expression of mGlu4 in the VP is high, circumscribed to afferent terminals, but may not be limited to D2-SPN terminals [43,44,106]. It might include some D1-SPN terminals and afferences from the thalamus (dorsomedial and midline nuclei) and the basolateral amygdala, which may have contributed to the beneficial effects of optogluram [43,44,76,106–108]. However, mGlu4-positive terminals detected in the VP form symmetrical synapses that are considered as GABAergic synapses [43]. This would exclude the participation of mGlu4 located at cortical, thalamic and basolateral amygdala afferent terminals from a participation to the effects of optogluram. Nonetheless, further investigations will be required to tease out the contribution of mGlu4 expressed on NAc D2-SPN terminals only. Importantly, photopharmacological experiments highlight the involvement of VP in controlling social interaction in *Fmr1^-/-^*mice. In this region, a finely tuned balance of activity between GABAergic and glutamatergic neuronal populations controls approach and avoidance behaviors, including social behaviors [109–112], in tight connection to the NAc D1/D2-SPN balance [32]. Interestingly, social avoidance triggered by optogenetic stimulation of D2-SPNs in mice was associated with increased *Fos* expression in the VP [30], suggesting increased neuronal activity consistent with previous studies [32]. Moreover, the relieving effects of mGlu4 PAM treatment in mice bearing NAc D1-SPN ablation on social behavior were associated with marked transcriptional downregulation in the VP [30]. Together, these results identify the VP, a major node of the basal ganglia circuit, as playing a crucial role in modulating social behavior in mice.

In conclusion, our study reveals, under ASD-relevant behavioral conditions, a bias in the D1/D2-SPN balance of activity towards excessive D2-SPN activity in *Fmr1^-/-^*mice, notably in the NAc [39,40]. This work also pinpoints the crucial contribution of VP, which integrates complex motivational information within the basal ganglia [107,109], in controlling social behavior. Furthermore, it demonstrates the therapeutic potential of drugs developed to repress D2-SPN activity to relieve core symptoms and co-morbidities of ASD, but also hyperactivity and cognitive deficits, in FXS.

## Acknowledgments

We acknowledge the technical assistance of Drs. Jorge Gandia and Thibaut Laboute, Yannick Corde, Audrey Léauté and Déborah Jaccaz in performing experiments. We thank the Experimental Unit PAO-1297 (EU0028, Animal Physiology Experimental Facility, DOI: 10.15454/1.5573896321728955E12) from the INRAe-Val de Loire Centre for animal breeding and care. Cell-based pharmacology assays were performed at the ARPEGE (Pharmacology Screening-Interactome) platform facility at the Institut de Génomique Fonctionnelle (Montpellier). We thank Drs. Xavier Rovira and Xavier Gomez-Santacana at MCS (IQAC-CSIC) and Drs. Carme Serra and Lourdes Muñoz at SimChem (IQAC-CSIC) for compound design and synthesis.

## Author contributions CRediT

**Mathieu Fonteneau:** Data Curation, Formal analysis, Investigation, Methodology, Visualization, Writing - Original Draft, Writing - Review & Editing. **Claire Terrier:** Conceptualization, Data Curation, Formal analysis, Investigation, Methodology, Writing - Original Draft, Writing - Review & Editing. **Florian Bolot:** Data Curation, Formal analysis, Visualization, Writing - Review & Editing. **Fani Pantouli:** Data Curation, Formal analysis, Investigation, Methodology, Writing - Review & Editing. **Fanny Malhaire:** Data Curation, Formal analysis, Investigation, Methodology, Writing - Review & Editing. **Roser Borràs-Tudurí:** Resources, Writing - Review & Editing. **Isabelle McCort:** Resources, Writing - Review & Editing. **Alexis Bailey:** Funding acquisition, Supervision, Writing - Review & Editing. **Francine Acher:** Funding acquisition, Resources, Supervision, Writing - Review & Editing. **Amadeu Llebaria:** Funding acquisition, Resources, Supervision, Writing - Review & Editing. **Cyril Goudet:** Conceptualization, Funding acquisition, Methodology, Supervision, Writing - Review & Editing. **Jérôme A.J. Becker:** Conceptualization, Data Curation, Formal analysis, Funding acquisition, Investigation, Methodology, Project administration, Supervision, Validation, Visualization, Writing - Original Draft, Writing - Review & Editing. **Julie Le Merrer:** Conceptualization, Data Curation, Formal analysis, Funding acquisition, Investigation, Methodology, Project administration, Supervision, Validation, Visualization, Writing - Original Draft, Writing - Review & Editing.

