## Supplementary tables and figures for "Tilted striatofugal balance and beneficial effects of facilitating mGlu4 receptor activity in the Fmr1^-/-^ mouse model of Fragile X Syndrome"

| Refseq | Gene name | Gene title | Forward oligonucleotide | Reverse oligonucleotide |
|---|---|---|---|---|
| NM_009630 | adenosine A2a receptor | *Adora2a* | TCAGCCTCTTGGCTATTGCC | CTCAAACAGACAGGTCACCCG |
| NM_028755 | cyclic AMP-regulated phosphoprotein, 21 | *Arpp21* | TGAGGAGGAGAAGCTGGAAC | GGTGACTTTCCCCTCCAGAT |
| NM_009732 | arginine vasopressine | *Avp* | ACACTACGCTCTTCCGCTTGT | CACTGTCTCAGCTCCATGTCA |
| NM_010076 | dopamine receptor D1A | *Drd1* | AGATCGGGCATTTGGAGAG | GGATGCTGCCTCTTCTTCTG |
| NM_010077 | dopamine receptor D2 | *Drd2* | TGCCATTGTTCTTGGTGTGT | GTGAAGGCGCTGTAGAGGAC |
| NM_016976 | glutamate receptor, metabotropic 1 | *Grm1* | TTCCCACCAGTTGAGAAACC | AAAGCCCTGACTGCACAAGT |
| NM_001160353 | glutamate receptor, metabotropic 2 | *Grm2* | CTTGTAGCTATGCCCCGTGT | GACTGGAAGCACCTTTGCAT |
| NM_181850 | glutamate receptor, metabotropic 3 | *Grm3* | GTGGTCTTGGGCTGTTTGTT | TGCTTGCAGAGGACTGAGAA |
| NM_001013385 | glutamate receptor, metabotropic 4 | *Grm4* | CTTTCTCTGCTATGCCACCACC | TAGCTGATGCTCATGCCAAGCC |
| NM_001143834 | glutamate receptor, metabotropic 5 | *Grm5* | CCACTGACGACTTGACAGGTT | CAGTAACGAAGAGGGTGGCTA |
| NM_177328 | glutamate receptor, metabotropic 7 | *Grm7* | CCAAAAACTGTGCTGGGAAG | GCATGCATTCCAAAGATCAA |
| NM_008174 | glutamate receptor, metabotropic 8 | *Grm8* | TCCTGTCCCACACTGTTCAA | AAAATTGCATGCGTCAATCA |
| NM_010471 | hippocalcin | *Hpca* | CCGGAAAAGAGGACTGAGAA | ATGAACTCCTCCAGCGACAG |
| NM_010444 | nuclear receptor subfamily 4, group A, member 1 | *Nr4a1* | CAATGCTTCGTGTCAGCACT | TGGCGCTTTTCTGTACTGTG |
| NM_011013 | opioid receptor, mu | *Oprm1* | ATCCTCTCTGAAGCCAACCC | TTCCTGTCTTGGGCCATCAT |
| NM_011025 | oxytocin | *Oxt* | CTGCTTGGCTTACTGGCTCT | GGGAGACACTTGCGCATATC |
| NM_001081147 | oxytocin receptor | *Oxtr* | CTTAGGGCCAAAAGGTGTCA | GCAGGTTTCTATGCCCTCTG |
| NM_018863 | prodynorphin | *Pdyn* | TTTGGCAACGGAAAAGAATC | TAGCGTTTGGCCTGTTTTCT |
| NM_001002927 | preproenkephalin | *Penk* | ATGCAGATGAGGGAGACACC | GCTTCTGCAGCTCTTTTGCT |

##### Table S1. List of primers used for qRT-PCR

**Analyzed data:** behavioral parameters from chronic treatment experiments; except those from marble burying and three-chamber tests (not performed in the same cohort for istradefylline experiment)

**Model:** Sex ~ Intercept + Grooming + Circling + Shakes + Latency to feed + %Same arm returns + Number of nose contacts + Time spent in nose contact + Number of paw contacts + Time spent in paw contact + Grooming after social contact + Following

| Odds ratios | Variable | Estimate | 95% CI (profile likelihood) | Likelihood ratio | P value | P value summary |
|---|---|---|---|---|---|---|
| β0 | Intercept | 2.764 | 0.6269 to 12.66 | 1.8 | 0.1797 | ns |
| β1 | Grooming | 1.01 | 0.7823 to 1.301 | 0.00586 | 0.939 | ns |
| β2 | Circling | 0.9012 | 0.7588 to 1.065 | 1.485 | 0.223 | ns |
| β3 | Shakes | 1.05 | 0.8949 to 1.234 | 0.3571 | 0.5501 | ns |
| β4 | Latency to feed | 1.003 | 0.9987 to 1.007 | 1.615 | 0.2038 | ns |
| β5 | %Same arm returns | 0.945 | 0.8758 to 1.013 | 2.511 | 0.1131 | ns |
| β6 | Number of nose contacts | 0.9852 | 0.9493 to 1.022 | 0.644 | 0.4223 | ns |
| β7 | Time spent in nose contact | 0.9973 | 0.9681 to 1.027 | 0.03152 | 0.8591 | ns |
| β8 | Number of paw contacts | 0.9578 | 0.5989 to 1.530 | 0.03312 | 0.8556 | ns |
| β9 | Time spent in paw contact | 1.006 | 0.7121 to 1.409 | 0.001255 | 0.9717 | ns |
| β10 | Grooming after social contact | 0.9268 | 0.6881 to 1.244 | 0.2583 | 0.6113 | ns |
| β11 | Following | 1.005 | 0.8843 to 1.140 | 0.004927 | 0.944 | ns |
| **Pseudo R squared** | | | | | | |
| Tjur's R squared | 0.0402 | | | | | |
| Nagelkerke's R squared | 0.05373 | | | | | |
| **Hypothesis tests** | **Statistic** | **P value** | **Null hypothesis** | **Reject Null Hypothesis?** | **P value summary** | |
| Log-likelihood ratio (G squared) | 7.361 | 0.7692 | Simpler (intercept-only) model is correct | No | ns | |
| **Data summary** | | | | | | |
| Rows in table | 187 | | | | | |
| Rows skipped (missing data) | 8 | | | | | |
| Rows analyzed (#observations) | 179 | | | | | |
| Number of F | 87 | | | | | |
| Number of M | 92 | | | | | |
| Number of parameter estimates | 12 | | | | | |
| #observations/#parameters | 14.9 | | | | | |
| # of F/#parameters | 7.3 | | | | | |
| # of M/#parameters | 7.7 | | | | | |

##### Table S2. Multiple logistic regression between behavior and se

**Analyzed data:** principal component scores (PC1 and PC2) from the principal component analysis of behavioral parameters (see Figure S1)
**Model:** Sex ~ Intercept + PC1 + PC2

| Odds ratios | Variable | Estimate | 95% CI (profile likelihood) | Likelihood ratio | P value | P value summary |
|---|---|---|---|---|---|---|
| β0 | Intercept | 0.9452 | 0.7039 to 1.268 | 0.1415 | 0.7068 | ns |
| β1 | PC1 | 1.001 | 0.8726 to 1.150 | 0.0003492 | 0.9851 | ns |
| β2 | PC2 | 0.8956 | 0.7067 to 1.127 | 0.8787 | 0.3485 | ns |

| Pseudo R squared | |
|---|---|
| Tjur's R squared | 0.004854 |
| Nagelkerke's R squared | 0.006533 |

| Hypothesis tests | Statistic | P value | Null hypothesis | Reject Null Hypothesis? | P value summary |
|---|---|---|---|---|---|
| Log-likelihood ratio (G squared) | 0.879 | 0.6444 | Simpler (intercept-only) model is correct | No | ns |

| Data summary | |
|---|---|
| Rows in table | 187 |
| Rows skipped (missing data) | 8 |
| Rows analyzed (#observations) | 179 |
| Number of F | 87 |
| Number of M | 92 |
| Number of parameter estimates | 3 |
| #observations/#parameters | 59.7 |
| # of F/#parameters | 29 |
| # of M/#parameters | 30.7 |

##### Table S3. Principal component logistic regression between behavior and sex

### Supplementary figures

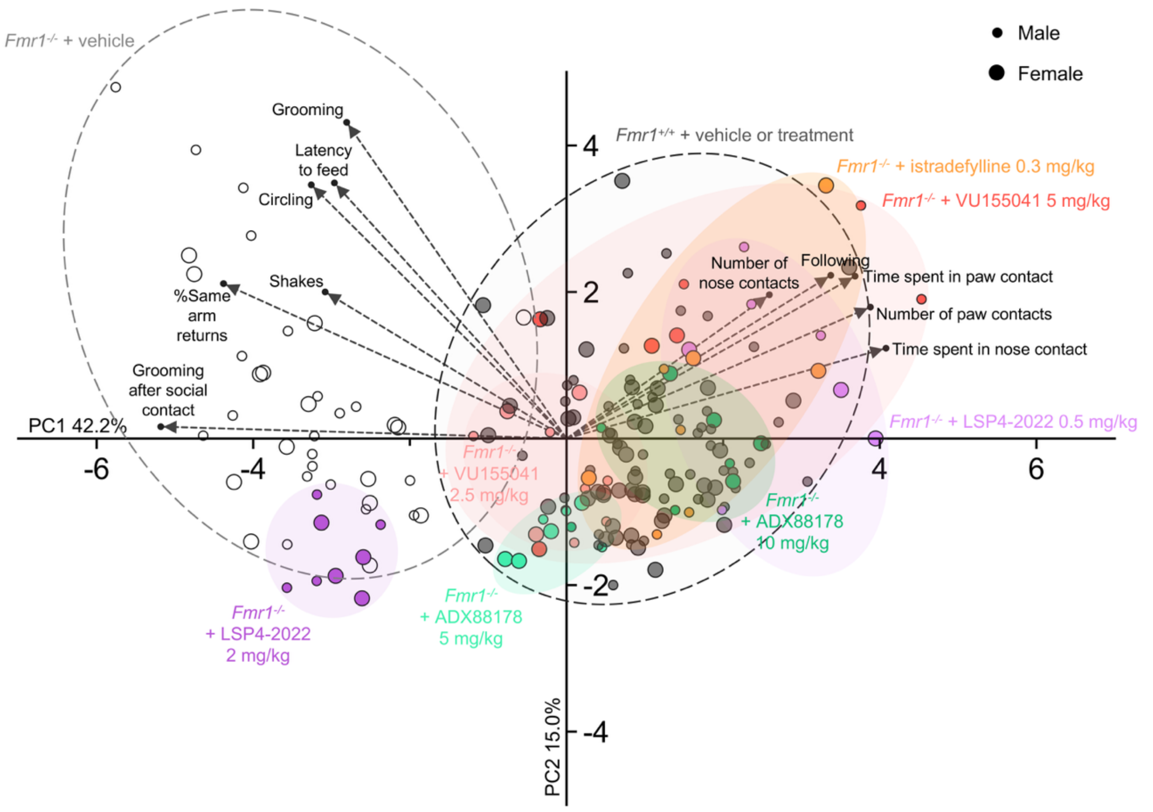


**Figure S1: Principal component analysis of ASD-like behavioral responses in *Fmr1*$^{+/+}$ and *Fmr1*$^{-/-}$ mice treated with vehicle or pharmacological compounds.** To evaluate the influence of sex on ASD-like behavior in *Fmr1*$^{-/-}$ and *Fmr1*$^{+/+}$ mice, we performed a principal component analysis using behavioral parameters longitudinally measured in the same mice during chronic treatment experiments; except from marble burying and three-chamber tests that were not performed in the same cohort for the istradefylline experiment. All the cohorts that undergone chronic pharmacological treatment were used for this analysis: mGlu4 PAM VU0155041 (0, 2.5 and 5 mg/kg) and ADX88178 (0, 5 and 10 mg/kg), mGlu4 agonist: LSP4-2022 (0, 0.5 and 2 mg/kg) and A2A antagonist: istradefylline (0 and 0.3 mg/kg). The analysis segregated behavioral parameters along two principal components (PCs) with PC1 showing opposed pro-social and ASD-like behaviors. Male and female individuals were evenly distributed, suggesting that sex had little influence on behavior in *Fmr1*$^{-/-}$ mice under different treatment conditions. This absence of effect was confirmed by multiple logistic regression between sex and behavioral data (Table S2) and by logistic regression between sex, PC1 and PC2 (Table S3). In contrast with sex, treatment and genotype had a major effect on subject distribution. *Fmr1*$^{-/-}$ mice under mGlu4 PAM, mGlu4 agonist (low dose) or A2A antagonist treatment overlapped with *Fmr1*$^{+/+}$ mice (vehicle and compound-treated), illustrating a normalization of their behavior. In contrast, *Fmr1*$^{-/-}$ mice treated with the mGlu4 agonist LSP4-2022 at 2 mg/kg overlapped with vehicle-treated *Fmr1*$^{-/-}$ mice, illustrating the absence of beneficial effect for this dose on ASD-like behaviors.

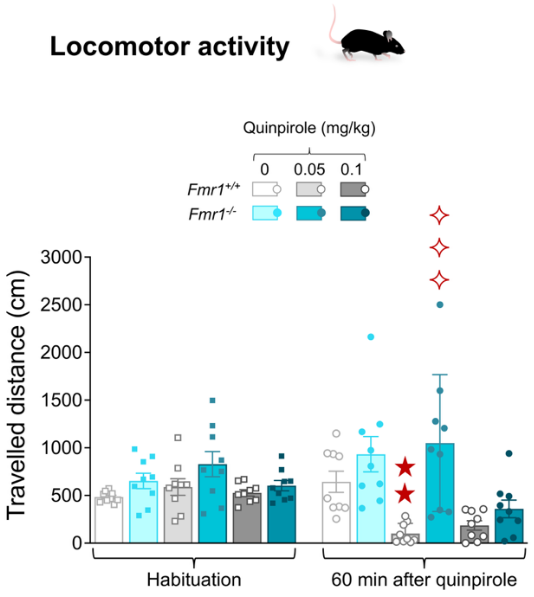


**Figure S2. Effect on quinpirole on locomotor activity in *Fmr1*$^{-/-}$ mice.** Adult *Fmr1*$^{+/+}$ and *Fmr1*$^{-/-}$ mice received the D2/D3-type receptor agonist quinpirole (0, 0.05 and 0.1 mg/kg, s.c.; n=9 mice per group) after a 15-min habituation period. Over a one-hour period, quinpirole decreased the locomotion of *Fmr1*$^{+/+}$ mice treated with 0.05 mg/kg (travelled distance in 60 min: $H_{5,54}=32.2$, $p<0.001$). However, unlike during the first 30 minutes after injection (Figure 1), this effect was no longer noticeable in *Fmr1*$^{+/+}$ mice treated with 0.1 mg/kg. In *Fmr1*$^{-/-}$ mice, quinpirole failed to decrease locomotor activity. Results are shown as scatter plots and mean ± SEM. Solid stars: comparison with the vehicle-treated *Fmr1*$^{+/+}$ group; diamonds: comparison with the *Fmr1*$^{+/+}$ group treated with the same dose of treatment. Two symbols: p<0.01; three symbols: p<0.001.

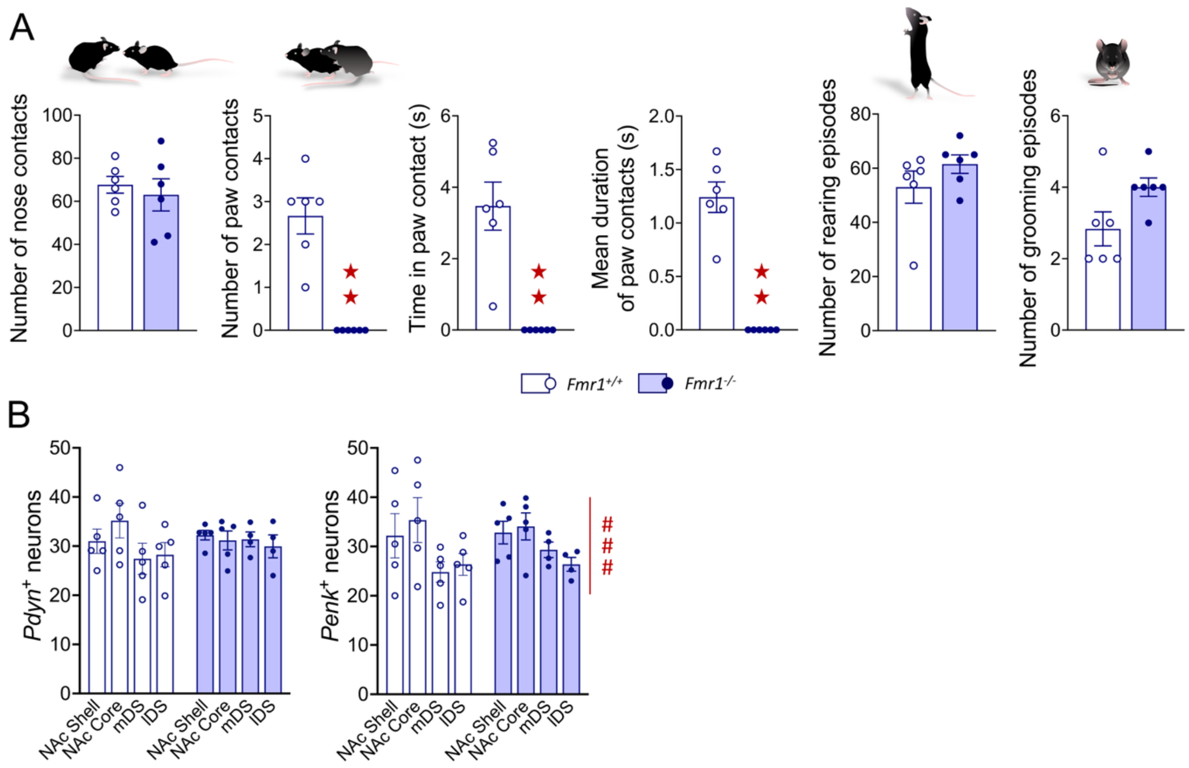


**Figure S3. Assessment of *Fos* positive of D1- and D2-SPNs following direct social interaction in *Fmr1*$^{-/-}$ mice using RNAscope® hybridization. (A)** *Fmr1*$^{+/+}$ and *Fmr1*$^{-/-}$ mice displayed similar numbers of nose contacts, rearing episodes (vertical activity) and grooming episodes; in contrast, *Fmr1*$^{-/-}$ mice displayed a severe deficit in the number ($U=0.0$, $p<0.01$), time spent in ($U=0.0$, $p<0.01$) and mean duration of paw contacts ($U=0.0$, $p<0.01$). **(B)** The number of *Pdyn*-positive and *Penk*-positive neurons were similar between *Fmr1*$^{+/+}$ and *Fmr1*$^{-/-}$ mice; the number of *Penk*-positive neurons was lower in dorsal compared to ventral areas of the striatum, in both mouse lines ($R$: $F_{3,20}=16.6$, $p<0.001$). Behavioral and hybridization data are shown as scatter plots and mean ± SEM. Solid stars: comparison with the *Fmr1*$^{+/+}$ group; hashtag: region effect. One symbol: p<0.05; two symbols: p<0.01. Abbreviations: lDS: dorsal striatum, lateral part; mDS: dorsal striatum, medial part; NAc Core: core of the nucleus accumbens; NAc Shell: shell of the nucleus accumbens; R: region effect.

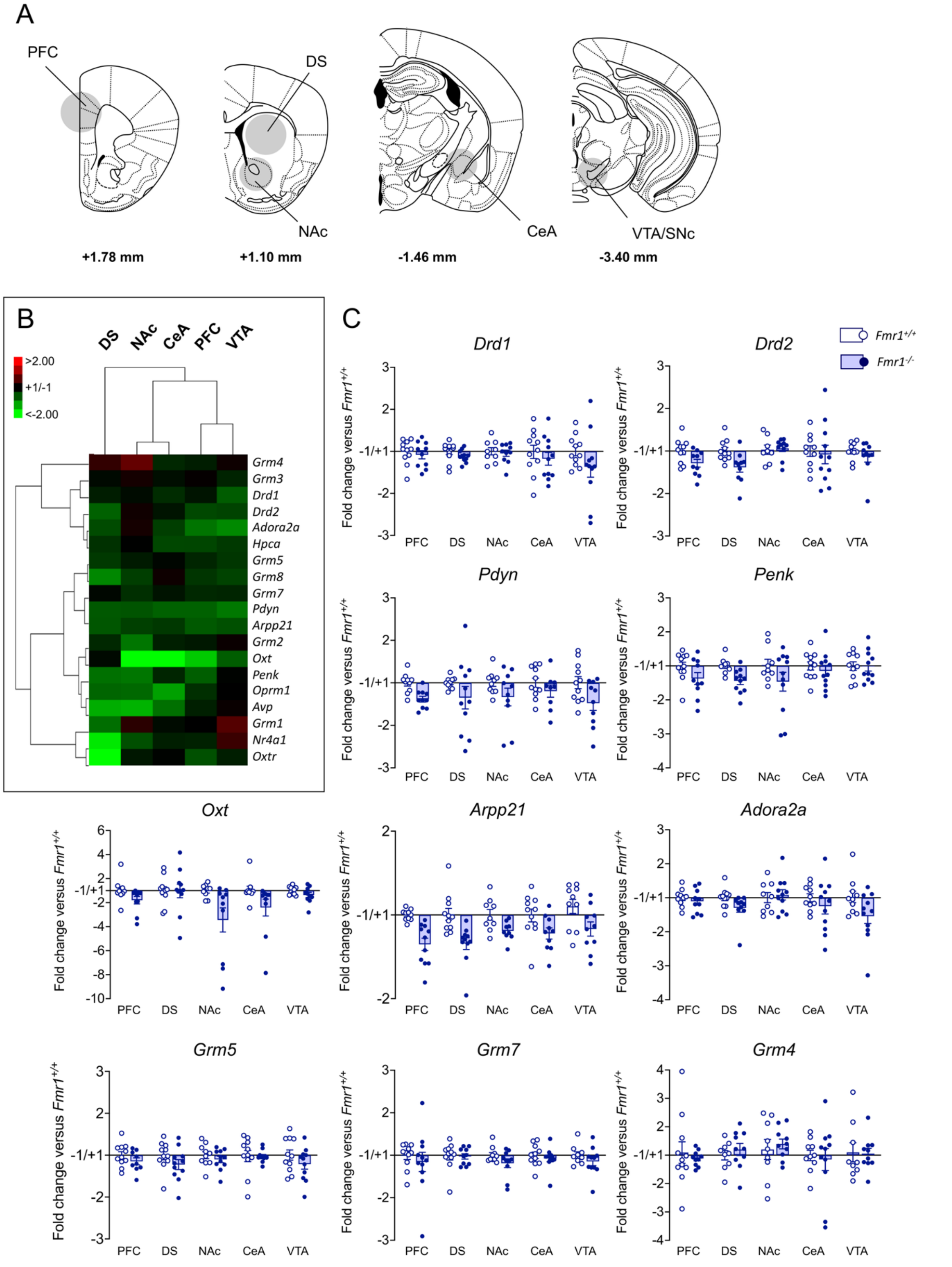


**Figure S4. Transcriptional effects of *Fmr1* deletion for a collection of genes in five brain regions. (A)** Schematic representations depict brain regions dissected for gene expression study. PFC (one medial punch ⦰ 1.25 mm), DS (one punch/side, ⦰ 2 mm), NAc, CeA, VTA/SNc (one punch/side, ⦰ 1.25 mm) were punched on 1-mm thick brain slices. Coordinates refer to bregma. **(B)** Clustering analysis reveals a global downregulation of gene expression in *Fmr1*$^{-/-}$

mice. **(C)** The expression of markers of D1-SPNs (*Pdyn*, *G:* $F_{1,92}$*=13.6, p<0.001*), D2-SPNs (*Penk*, *G:* $F_{1,96}$*=8.6, p<0.01*; *Adora2a*, *G:* $F_{1,93}$*=5.3, p<0.05*) or both SPNs types (*Arpp21*, *G:* $F_{1,20}$*=35.7, p<0.001*) was decreased in *Fmr1*$^{-/-}$ mice, as well as the expression of the neuropeptide oxytocin (*Oxt*, *G:* $F_{1,20}$*=6.1, p<0.05*). In contrast, no such significant downregulation was detected for *Drd1*, *Drd2*, *Grm4*, *Grm5* and *Grm7*. Transcriptional results (fold change compared to *Fmr1*$^{+/+}$ mice) are shown as scatter plots and mean ± SEM. Abbreviations: CeA: central amygdala; DS: dorsal striatum; G: genotype effect; NAc: nucleus accumbens; PFC: prefrontal cortex; VTA/SNc: ventral tegmental area/substantia nigra, pars compacta.

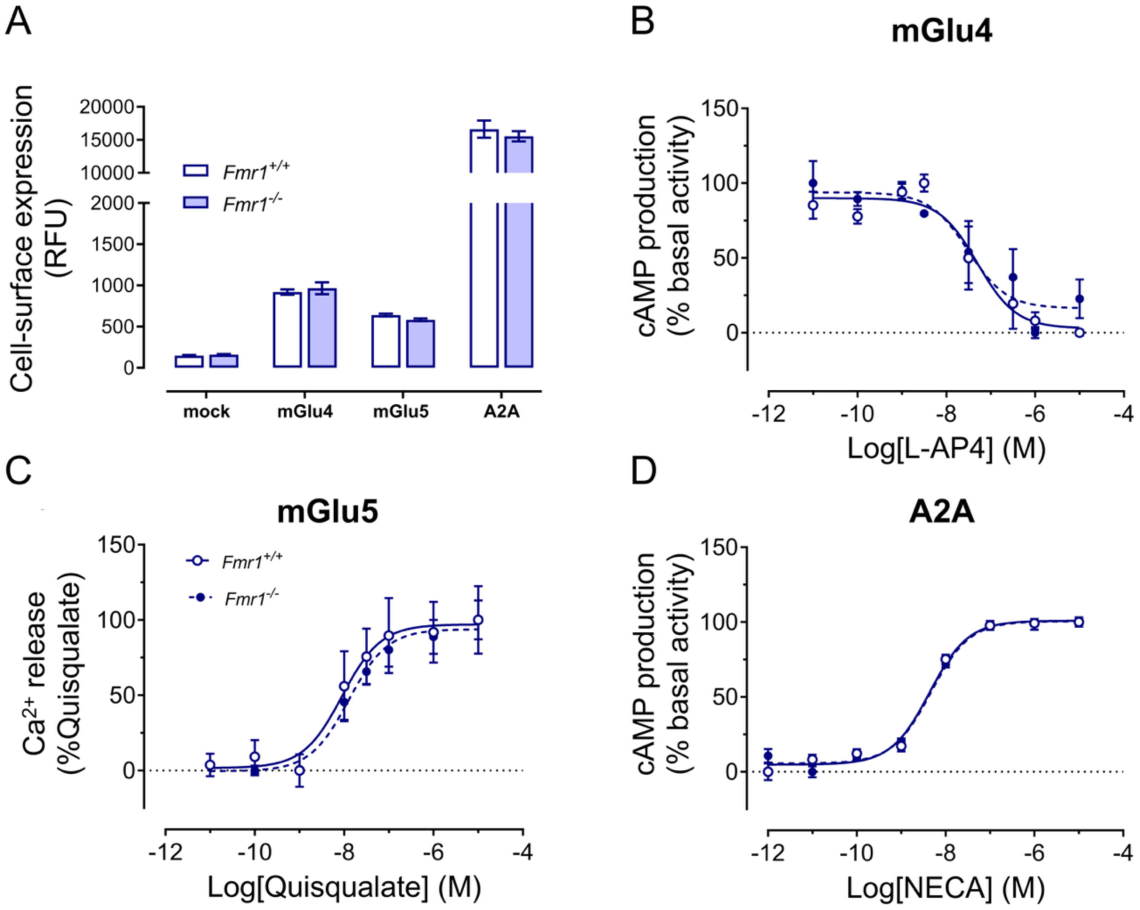


**Figure S5. The lack of *Fmr1* does not impair the coupling of mGlu4, mGlu5 and A2A receptors in heterologous HEK293 cells.** SNAP-tagged human mGlu4, mGlu5 or A2A receptors were transiently transfected in *Fmr1*$^{-/-}$ or *Fmr1*$^{+/+}$ HEK293 cells, then their cell surface expression and their pharmacological response were compared 24 h after transfection. **(A)** Cell surface expression of mGlu4, mGlu5 or A2A does not differ between *Fmr1*$^{-/-}$ and *Fmr1*$^{+/+}$ HEK293 cells, as determined following labelling by SNAP-Lumi4-Tb. Results are shown as column bars and mean ± SEM (mGlu4: n=8 independent experiments; mGlu5: n=3 independent experiments; A2A: n=4 independent experiments). **(B)** Dose-dependent inhibition of Forskolin (300 nM) evoked cAMP production following stimulation of mGlu4 receptors by L-

AP4, as determined by the Cisbio-bioassays $G_i$ cAMP kit. The potency (pEC50) of the mGlu4 reference agonist is not modified between *Fmr1*$^{+/+}$ and *Fmr1*$^{-/-}$ cells: 7.30 ± 0.25 vs 7.51 ± 0.31. Curves represent the mean ± SEM of n=4 independent experiments. **(C)** Dose-dependent release of intracellular $Ca^{2+}$ following stimulation of mGlu5 receptors by quisqualate, as determined by the fluorescent $Ca^{2+}$ sensitive dye Fluo4AM. pEC50 of the mGlu5 reference agonist is not modified between *Fmr1*$^{+/+}$ and *Fmr1*$^{-/-}$ cells: 8.05 ± 0.32 vs 7.90 ± 0.15. Curves represent the mean ± SEM of n=5 independent experiments. **(D)** Dose-dependent production of cAMP following stimulation of A2A receptors by NECA, as determined by the Cisbio-bioassays $G_s$ cAMP kit. pEC50 of the A2A reference agonist is not modified between *Fmr1*$^{+/+}$ and *Fmr1*$^{-/-}$ cells: 8.36 ± 0.07 vs 8.33 ± 0.07. Curves represent the mean ± SEM of n=4 independent experiments.

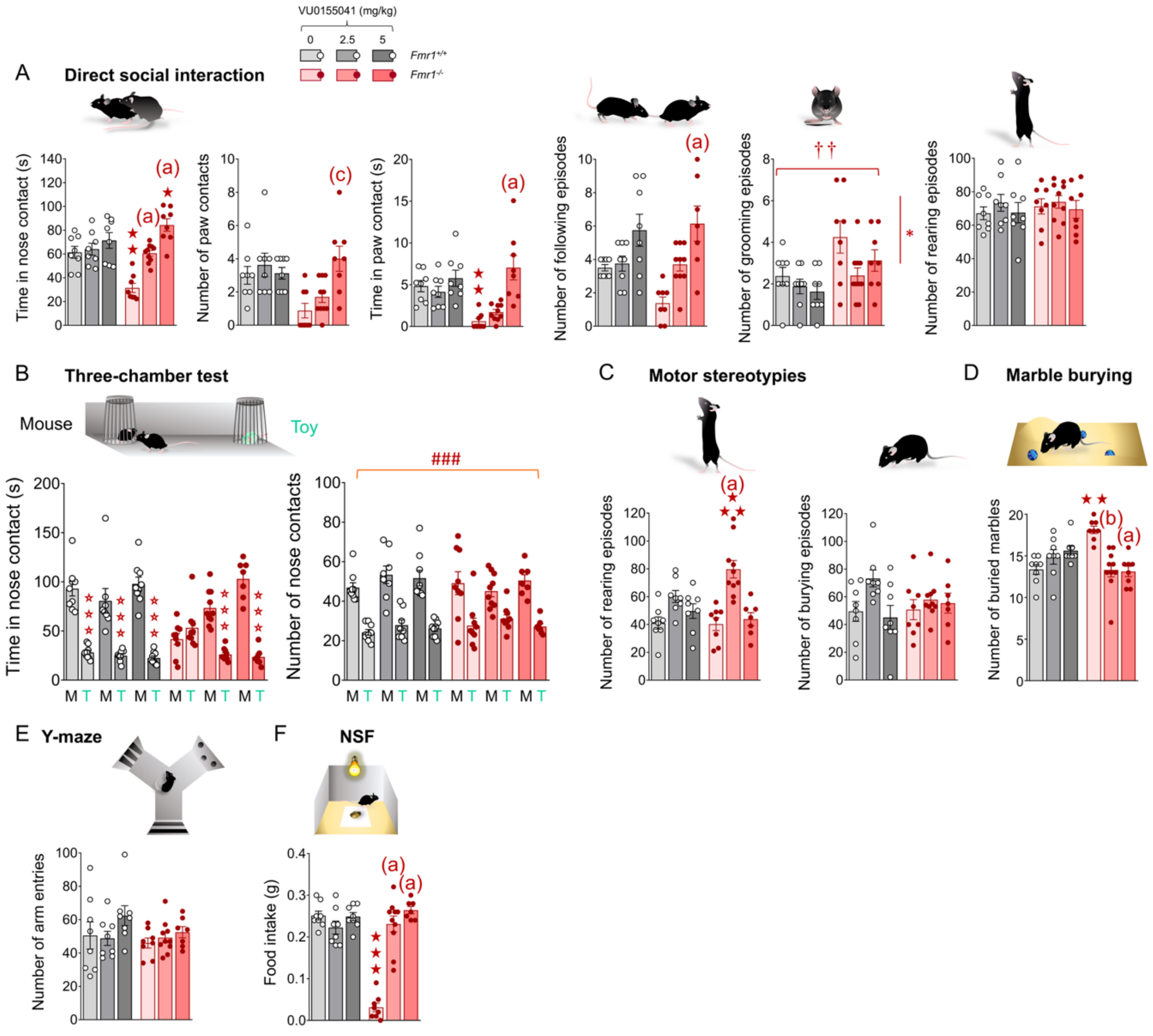


**Figure S6. Effects of chronic facilitation of mGlu4 activity on ASD-like behavioral responses in *Fmr1*$^{-/-}$ mice. (A)** In the direct social interaction test, chronic VU0155041 administration dose-dependently normalized the time spent in nose contact (*G x Tr:* $F_{2,44}=10.1$, $p<0.001$), the number ($H_{5,50}=19.1$, $p<0.01$) and time spent in paw contacts ($H_{5,50}=29.9$,

$p<0.001$), as well as the number of following episodes ($H_{5,50}=22.9, p<0.001$) in *Fmr1*$^{-/-}$ mice. The total number of grooming episodes was higher in *Fmr1*$^{-/-}$ mice, and VU0155041 decreased this number in both mouse lines ($G: F_{1,44}=10.1, p<0.01; Tr: F_{2,44}=3.3, p<0.05$). The number of rearing episodes (vertical activity) was not affected. **(B)** In the three-chamber test, chronic VU0155041 treatment normalized the time spent by *Fmr1*$^{-/-}$ mice in contact with the mouse versus the toy ($G \times Tr \times S: F_{2,46}=10.5, p<0.001$); mice made globally more nose contacts with the mouse than with the toy ($S: F_{1,46}=102.4, p<0.001$). **(C)** *Fmr1*$^{-/-}$ mice treated with VU0155041 at 2.5 mg/kg displayed more rearing episodes during motor stereotypies assessment ($G \times Tr: F_{2,43}=3.5, p<0.05$). This treatment had no effect on burying behavior. **(D)** In the marble burying test, VU0155041 normalized the number of buried marbles at both doses ($H_{5,50}=25.4, p<0.001$). **(E)** In the Y-maze, VU0155041 had no influence on the number of arm entries. **(F)** In the novelty-suppressed feeding test, VU0155041 restored food intake in *Fmr1*$^{-/-}$ mice ($G \times Tr: F_{2,43}=46.1, p<0.001$). Results are shown as scatter plots and mean ± SEM. Solid stars: comparison with the vehicle-treated *Fmr1*$^{+/+}$ group; open stars: genotype x treatment x stimulus interaction, mouse versus toy comparison; hashtags: stimulus effect; daggers: genotype effect; asterisks: treatment effect. One symbol: p<0.05; two symbols: p<0.01; three symbols: p<0.001. Letters: comparison with vehicle-treated *Fmr1*$^{-/-}$ group; (c): p<0.05, (b): p<0.01, (a): p<0.001. Abbreviations: G: genotype effect; M: mouse; NSF: novelty-suppressed feeding test; S: stimulus effect; Tr: treatment effect; T: toy.

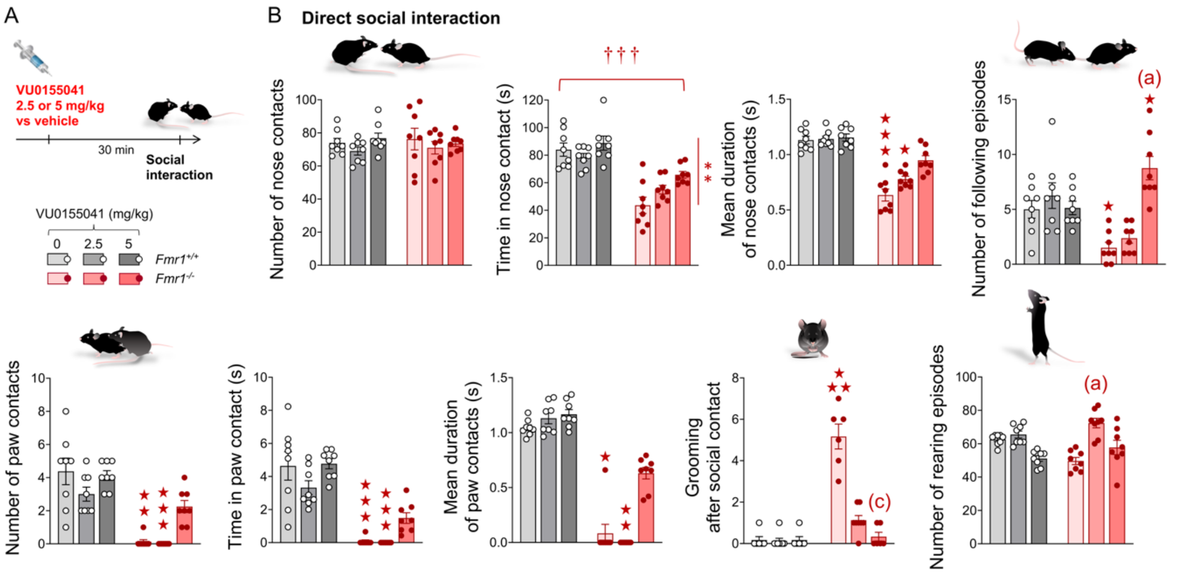


**Figure S7. Effects of a single administration of VU0155041 on direct social interaction in *Fmr1*$^{-/-}$ mice.** When administered once 30 min before testing, VU0155041 increased the time spent in nose contact in all mice ($Tr: F_{2,42}=5.6, p<0.01$) but failed to completely restore this parameter in *Fmr1*$^{-/-}$ mice ($G: F_{1,42}=70.2, p<0.001$); at the dose of 5 mg/kg, it partially restored the duration of nose contacts in *Fmr1*$^{-/-}$ mice ($H_{5,48}=39.6, p<0.001$), the number of ($H_{5,48}=36.7, p<0.001$), time spent in ($H_{5,48}=39.3, p<0.001$) and mean duration of ($H_{5,48}=41.1,$

*$p<0.001$*) paw contacts. This highest dose normalizes the number of following episodes (*G x Tr: $F_{2,42}=13.5$, $p<0.001$*) and suppressed grooming after social contact (*$H_{5,48}=33.2$, $p<0.001$*) in *Fmr1*$^{-/-}$ mice. At 2.5 mg/kg, acute VU0155041 administration increased the number of rearing episodes (*$H_{5,48}=29.9$, $p<0.001$*) in *Fmr1*$^{-/-}$ mice. Results are shown as scatter plots and mean ± SEM. Solid stars: comparison with the vehicle-treated *Fmr1*$^{+/+}$ group; daggers: genotype effect. One symbol: $p<0.05$; two symbols: $p<0.01$; three symbols: $p<0.001$. Letters: comparison with vehicle-treated *Fmr1*$^{-/-}$ group; (c): $p<0.05$, (a): $p<0.001$. Abbreviations: G: genotype effect; Tr: treatment effect.

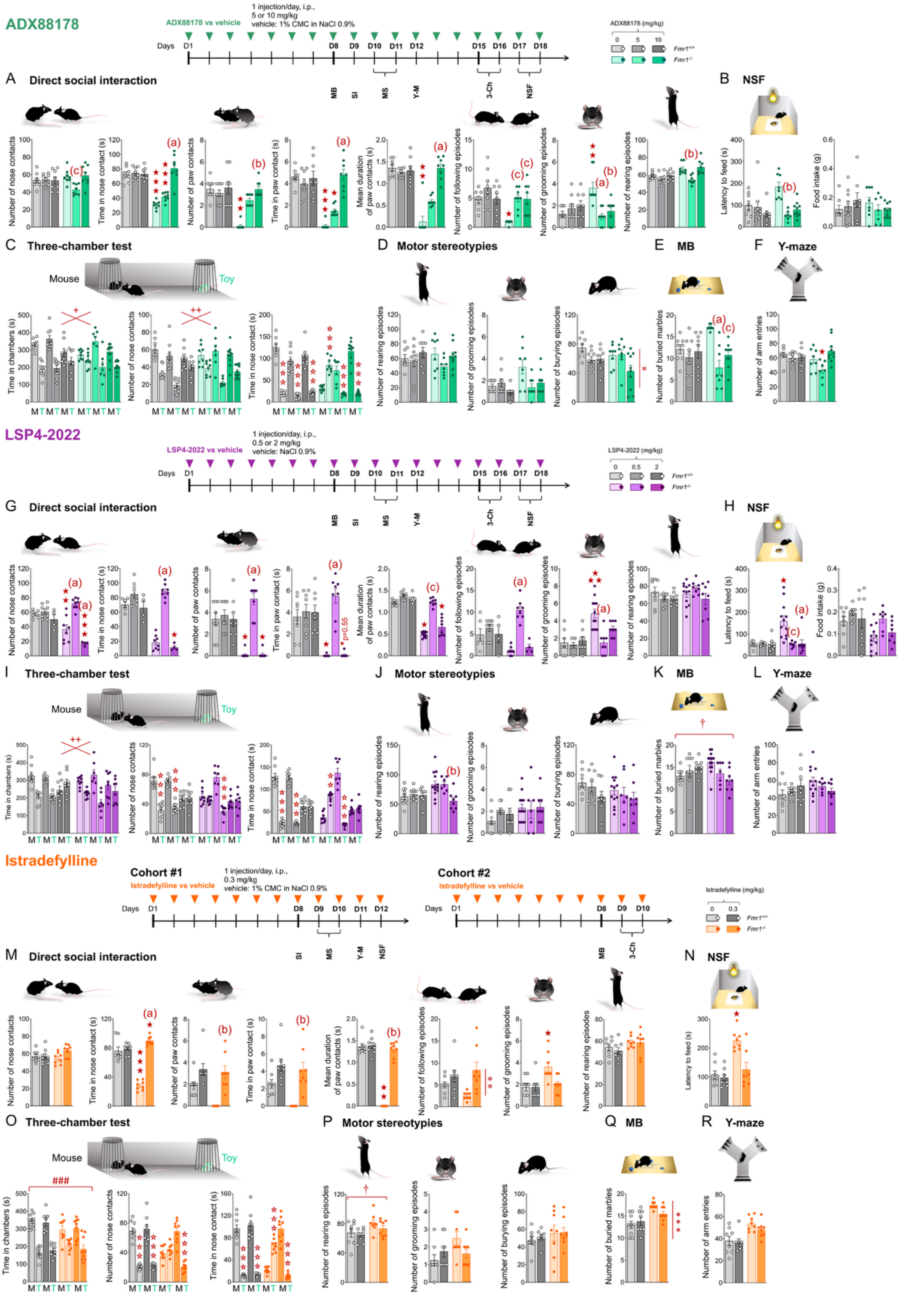

**Figure S8. Compared effects of chronic mGlu4 facilitation or orthosteric activation, and adenosine A2A receptor blockade on core ASD-like symptoms in *Fmr1*$^{-/-}$ mice.** A first cohort of *Fmr1*$^{+/+}$ and *Fmr1*$^{-/-}$ mice (n=8-10 per group) was injected once daily with either the mGlu4 PAM ADX88178 (5 or 10 mg/kg, i.p.) or vehicle (1% CMC in NaCl 0.9%) for 18 days. **(A)** In the direct social interaction test, *Fmr1*$^{-/-}$ mice treated with ADX88178 at 5 mg/kg made fewer nose contacts ($G \times Tr$: $F_{2,42}=4.5$, $p<0.05$) and rearing episodes ($H_{5,48}=22.2$, $p<0.001$); ADX88178 treatment dose-dependently restored the time spent in nose ($G \times Tr$: $F_{2,42}=25.4$, $p<0.001$) and paw contacts ($H_{5,48}=33.1$, $p<0.001$), the number of paw contacts ($H_{5,48}=25.2$, $p<0.001$) and their duration ($H_{5,48}=32.8$, $p<0.001$), as well as the number of following ($H_{5,48}=20.9$, $p<0.001$) and grooming episodes ($G \times Tr$: $F_{2,42}=9.2$, $p<0.001$) in *Fmr1*$^{-/-}$ mice. **(B)** In the NSF, ADX88178 at 5 mg/kg reduced the latency to feed in *Fmr1*$^{-/-}$ mice ($H_{5,50}=18.8$, $p<0.01$). **(C)** In the three-chamber test, ADX88178 globally increased the time spent in the chamber with the mouse ($Tr \times S$: $F_{2,44}=3.5$, $p<0.05$) and the number of nose contacts with the mouse ($Tr \times S$: $F_{2,44}=6.1$, $p<0.01$). This treatment fully rescued the preference for spending more time with the mouse over the toy in *Fmr1*$^{-/-}$ mice ($G \times Tr \times S$: $F_{2,44}=69.4$, $p<0.001$). **(D)** ADX88178 had no significant effects on rearing and grooming behaviors but generally decreases the number of burying episodes ($Tr$: $F_{2,45}=4.6$, $p<0.05$) during the assessment of motor stereotypies. **(E)** ADX88178 decreased marble burying in *Fmr1*$^{-/-}$ mice ($H_{5,50}=22.1$, $p<0.001$). **(F)** In the Y-maze, ADX88178 at 5 mg/kg reduced the number of arm entries ($G \times Tr$: $F_{2,44}=4.1$, $p<0.05$).

A second cohort of *Fmr1*$^{+/+}$ and *Fmr1*$^{-/-}$ mice (n=8-13 per group) was injected once daily with either the mGlu4 orthosteric agonist LSP4-2022 (0.5 or 2 mg/kg, i.p.) or vehicle (NaCl 0.9%) for 18 days. **(G)** In the direct social interaction test, LSP4-2022 at the dose of 0.5 mg/kg, but not 2 mg/kg, normalized the number of ($G \times Tr$: $F_{2,46}=25.3$, $p<0.001$) and time spent in ($H_{5,52}=40.5$, $p<0.001$) nose contacts, the number of ($H_{5,52}=40.3$, $p<0.001$), time spent in ($H_{5,52}=39.8$, $p<0.001$) and mean duration of ($H_{5,52}=40.2$, $p<0.001$) paw contacts, as well as the number of following ($H_{5,52}=36.3$, $p<0.001$) and grooming ($G \times Tr$: $F_{2,46}=6.2$, $p<0.01$) episodes. This compound had no effect on rearing activity. **(H)** In the NSF, LSP4-2022 at both doses normalized the latency to feed ($H_{5,52}=23.1$, $p<0.001$). **(I)** In the three-chamber test, LSP4-2022 globally increased the time spent in the chamber with the mouse ($Tr \times S$: $F_{2,47}=7.6$, $p<0.01$). At the dose of 0.5 mg/kg, LSP4-2022 restored the preference for making more frequent ($G \times Tr \times S$: $F_{2,47}=9.8$, $p<0.001$) and time spent in ($G \times Tr \times S$: $F_{2,47}=52.4$, $p<0.001$) nose contacts with the mouse over the toy, while under LSP4-2022 at 2 mg/kg mice failed to show such a preference. **(J)** LSP4-2022 at 2 mg/kg decreased rearing ($G \times Tr$: $F_{2,47}=4.0$, $p<0.05$) during the assessment of motor stereotypies. **(K)** In the marble burying test, *Fmr1*$^{-/-}$ mice buried more marbles than *Fmr1*$^{+/+}$ mice ($G$: $F_{1,47}=6.4$, $p<0.05$). **(L)** In the Y-maze, LSP4-2022 had no effect on the number of arm entries.

A third cohort of *Fmr1*$^{+/+}$ and *Fmr1*$^{-/-}$ mice (n=8 per group) was injected once daily with either the A2A receptor antagonist istradefylline (0.3 mg/kg, i.p.) or vehicle (1% CMC in NaCl 0.9%) for 12 days. A fourth cohort (n=9 per group) received either istradefylline or vehicle for 10 days.

**(M)** In the direct social interaction test, istradefylline rescued the time spent in nose contact (*G x Tr:* $F_{1,28}$*=74.1, p<0.001*), the number of ($H_{3,32}$*=22.1, p<0.001*), time spent in ($H_{3,32}$*=21.0, p<0.001*) and mean duration of ($H_{3,32}$*=18.0, p<0.001*) paw contacts. This treatment increased the number of following episodes in all mice (*Tr:* $F_{1,28}$*=11.1, p<0.01*) and reduced the number of grooming episodes in *Fmr1*$^{-/-}$ mice ($H_{3,32}$*=11.4, p<0.01*). **(N)** In the NSF, istradefylline reduced the latency to feed in *Fmr1*$^{-/-}$ mice ($H_{3,32}$*=14.0, p<0.01*); food intake was not measured in this experiment. **(O)** In the three-chamber test, all mice spent longer in the chamber with the mouse (*S:* $F_{1,30}$*=62.9, p<0.001*); istradefylline restored a preference for making more frequent (*G x Tr x S:* $F_{1,30}$*=22.4, p<0.001*) and time spent in (*G x Tr x S:* $F_{1,30}$*=76.1, p<0.001*) nose contacts with the mouse over the toy in *Fmr1*$^{-/-}$ mice. **(P)** *Fmr1*$^{-/-}$ mice made more frequent rearing episodes (*G:* $F_{1,30}$*=6.3, p<0.05*) during the assessment of motor stereotypies. **(Q)** Istradefylline decreased the number of buried marbles (*Tr:* $F_{1,30}$*=20.2, p<0.001; G x Tr:* $F_{1,30}$*=3.9, p=0.056*). **(R)** Istradefylline had no effect on the number of arm entries in the Y-maze. Results are shown as scatter plots and mean ± SEM. Solid stars: comparison with the vehicle-treated *Fmr1*$^{+/+}$ group; open stars: genotype x treatment x stimulus interaction, mouse versus toy comparison; plus signs: stimulus x treatment interaction; asterisks: treatment effect; hashtags: stimulus effect; daggers: genotype effect. One symbol: p<0.05; two symbols: p<0.01; three symbols: p<0.001. Letters: comparison with vehicle-treated Fmr1$^{-/-}$ group; (c): p<0.05, (b): p<0.01, (a): p<0.001. Abbreviations: CMC: carboxymethyl cellulose; G: genotype effect; M: mouse; MB: marble burying test; NSF: novelty-suppressed feeding test; S: stimulus effect; Tr: treatment effect; T: toy.

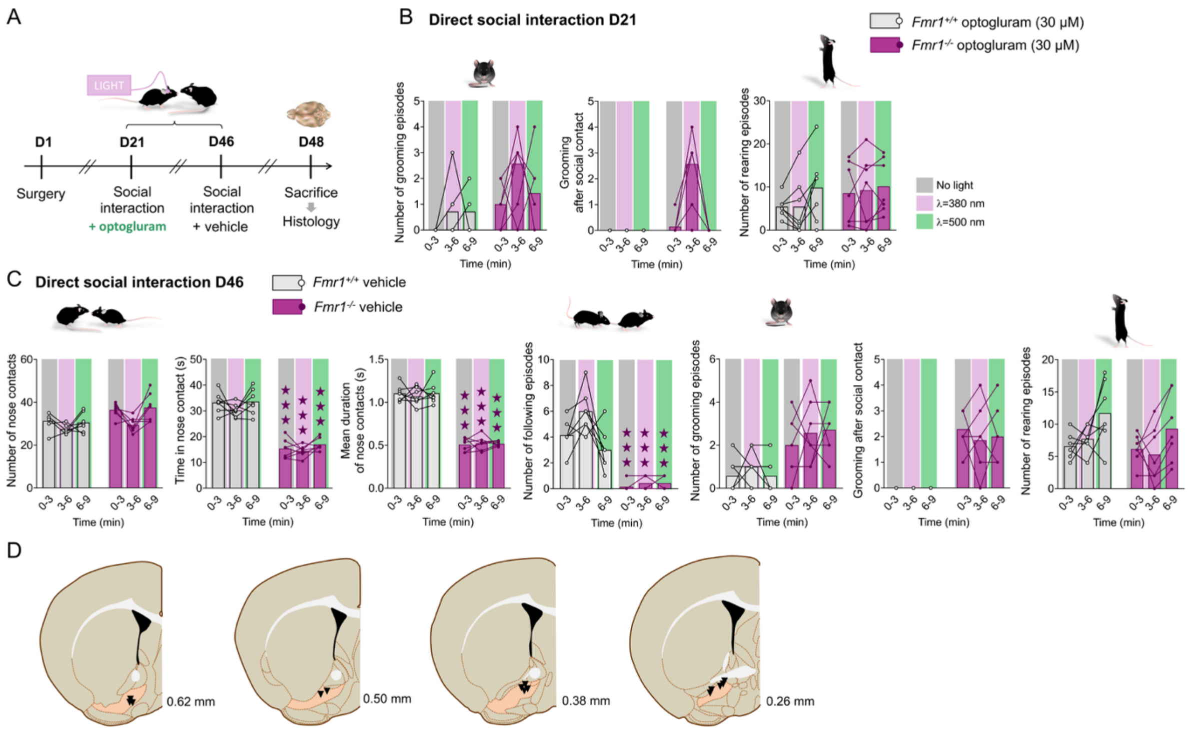


**Figure S9. Effects of photopharmacological activation within the ventral pallidum of the mGlu4 PAM optogluram on social interaction in *Fmr1*$^{-/-}$ mice. (A)** Timeline of the experiments. A first social interaction session was performed in the presence of optogluram at D21 following surgery (n=7 mice per group). A second session was performed at D46 during which the vehicle of optogluram (NaCl 0.9%) was microinjected into the VP. Mice were sacrificed at D48 for verification of cannula placement. **(B)** In the direct social interaction test (D21), the number of grooming episodes was increased in *Fmr1*$^{-/-}$ mice compared to *Fmr1*$^{+/+}$ mice (*G:* $F_{1,12}=29.8$, $p<0.001$). Switching optogluram on and off had no significant effect on the number of grooming events following a social contact or not, nor the number of rearing episodes (vertical activity). **(C)** On D46, the direct social interaction test was performed under vehicle microinjection. Under these conditions, *Fmr1*$^{-/-}$ mice displayed a severe deficit in time spent in nose contact, mean duration of nose contacts and number of following episodes. The number of grooming episodes was higher in *Fmr1*$^{-/-}$ mice compared to *Fmr1*$^{+/+}$ mice (*G:* $F_{1,12}=29.8$, $p<0.001$), while the number of rearing episodes was similar between genotypes. Illumination with violet or green light had no significant influence on these parameters. **(D)** Mapping of cannula placement on schematic brain sections for all mice. Coordinates refer to bregma. Results are shown as scatter plots and mean ± SEM. Solid stars: comparison with the optogluram-treated *Fmr1*$^{+/+}$ group under no light conditions; three symbols: p<0.001. Abbreviations: A: optogluram activation (on versus off); D: day; G: genotype; Tr: treatment (optogluram versus vehicle).
